# Follicle stimulating hormone signaling opposes the DRL-1/FLR-4 MAP Kinases to balance p38-mediated growth and lipid homeostasis in *C. elegans*

**DOI:** 10.1101/2023.01.07.523122

**Authors:** Sarah K. Torzone, Aaron Y. Park, Peter C. Breen, Natalie R. Cohen, Robert H. Dowen

## Abstract

Animals integrate developmental and nutritional signals before committing crucial resources to growth and reproduction; however, the pathways that perceive and respond to these inputs remain poorly understood. Here, we demonstrate that DRL-1 and FLR-4, which share similarity with mammalian mitogen-activated protein kinases, maintain lipid homeostasis in the *C. elegans* intestine. DRL-1 and FLR-4 function in a protein complex at the plasma membrane to promote development, as mutations in *drl-1* or *flr-4* confer slow growth, small body size, and impaired lipid homeostasis. To identify factors that oppose DRL-1/FLR-4, we performed a forward genetic screen for suppressors of the *drl-1* mutant phenotypes and identified mutations in *flr-2* and *fshr-1*, which encode the orthologues of follicle stimulating hormone and its putative G protein-coupled receptor, respectively. In the absence of DRL-1/FLR-4, neuronal FLR-2 acts through intestinal FSHR-1 and Protein Kinase A signaling to restrict growth. Furthermore, we show that opposing signaling through DRL-1 and FLR-2 coordinates TIR-1 phase transition, which modulates downstream p38/PMK-1 activity, lipid homeostasis, and development. Finally, we identify a surprising noncanonical role for the developmental transcription factor PHA-4/FOXA in the intestine where it restricts growth in response to impaired DRL-1 signaling. Our work uncovers a complex multi-tissue signaling network that converges on p38 signaling to maintain homeostasis during development.

## INTRODUCTION

Animals respond to environmental, nutritional, and developmental cues to balance resources between essential biological processes, ensuring fitness and reproductive fidelity. In metazoans, reproduction is a metabolically expensive process, requiring organisms to shift somatic energy stores to the germline to support the development of their offspring. This metabolic trade-off ensures reproductive fitness while restricting the somatic maintenance programs that support longevity (Kirkwood 1977; Kirkwood and Holliday 1979). The energetic balance between somatic and germline functions is coordinated by complex regulatory networks across diverse tissues that integrate developmental and environmental inputs; however, the homeostatic mechanisms that govern these metabolic trade-offs are not fully understood.

In many metazoans, including the nematode *Caenorhabditis elegans*, development into a reproductive adult is marked by production of vitellogenin proteins, which are structural and functional orthologues of the mammalian apoB protein that coordinates very low-density lipoprotein (VLDL) assembly, secretion, and reabsorption in the liver (Baker 1988). In *C. elegans*, the vitellogenins package intestinal lipids into VLDL-like particles, which are then secreted and captured by the LDL receptor RME-2 in oocytes (Kimble and Sharrock 1983; Grant and Hirsh 1999). The vitellogenin-associated lipids promote the recruitment of sperm to the oocyte during fertilization (Kubagawa et al. 2006), support robust development of the progeny (Perez et al. 2017), and facilitate larval survival during starvation conditions (Chotard et al. 2010; Van Rompay et al. 2015). While crucial for reproduction and the developmental success of the progeny, reallocation of these key lipid resources restricts maternal longevity (Seah et al. 2016; Ezcurra et al. 2018). Consistently, this metabolic trade-off can be finely tuned and is highly regulated by developmental, nutritional, and metabolic regulatory pathways (Murphy et al. 2003; DePina et al. 2011; Dowen et al. 2016; Goszczynski et al. 2016; Seah et al. 2016; Dowen 2019). The molecular basis of how these developmental regulators impact metabolic decisions to maintain organismal homeostasis is poorly understood.

Genetic screens aimed at uncovering genes required for the initiation of vitellogenesis have identified proteins with broader roles in development, metabolism, stress responses, and longevity (Van Rompay et al. 2015; Dowen et al. 2016; Dowen 2019). We previously identified the <u>d</u>ietary-<u>r</u>estriction-like gene *drl-1* in an RNAi screen as a candidate regulator of vitellogenesis (Dowen et al. 2016). DRL-1 is a serine-threonine mitogen activated protein kinase (MAPK) orthologous to mammalian MEKK3 that has been implicated in regulating metabolic, detoxification, and aging pathways (Chamoli et al. 2014, 2020). Loss of *drl-1* increases lifespan and upregulates detoxication genes, which requires the p38 MAPK signaling pathway (NSY-1/SEK-1/PMK-1); however, the dietary restriction-like metabolic state triggered by *drl-1* knockdown is not entirely dependent on p38 signaling (Chamoli et al. 2020). Interestingly, loss of a closely related MAP kinase gene, *flr-4*, induces a similar p38-dependent lifespan extension and induction of detoxication genes (Take-uchi et al. 2005; Verma et al. 2018). This observation suggests that DRL-1 and FLR-4 may function in the same signaling pathway; however, a biochemical association between these two proteins has not been demonstrated.

While the role of the p38/PMK-1 pathway in regulating innate immunity and oxidative stress responses is well defined (Kim et al. 2002; Aballay et al. 2003; Inoue et al. 2005), its function in development, as well as the molecular pathways that converge on p38 signaling to coordinate growth, are poorly understood. Here, we find that mutations in *drl-1* or *flr-4* severely impair development, growth, and lipid homeostasis in *C. elegans*, in part by governing the phase transition and activation of TIR-1/SARM1, a Toll/interleukin-1 receptor (TIR) domain-containing protein that activates p38/PMK-1 signaling (Peterson et al. 2022). We show that DRL-1 and FLR-4 are opposed by follicle stimulating hormone (FSH) signaling, which is mediated by the secreted neurohormone FLR-2, its putative intestinal G-protein coupled receptor FSHR-1, and downstream cAMP/Protein Kinase A (PKA) signaling. Moreover, our data suggest that these opposing pathways may converge on TIR-1 in the intestine to modulate p38 signaling and govern the subcellular localization of the PHA-4/FOXA transcription factor, a well-established regulator of development, to control growth and metabolic homeostasis. Thus, we demonstrate that intestinal p38/PMK-1 activity is coordinated by a non-cell-autonomous hormonal signal and intestinal MAPK pathway to maintain metabolic homeostasis and ensure robust development.

## RESULTS

### The DRL-1 protein kinase functions in the intestine to modulate fat transport and growth

The vitellogenin genes (*vit-1* through *vit-6*) are specifically expressed in the *C. elegans* intestine during the L4 larval to adult transition, coinciding with the onset of reproduction. This metabolic commitment can be precisely followed using a multi-copy P*vit-3::GFP* or a single-copy P*vit-3::mCherry* reporter transgene comprised of GFP or mCherry under the control of the *vit-3* promoter, respectively (Fig. 1A, S1A). Notably, the single-copy P*vit-3::mCherry* transgene is highly sensitive and its expression closely resembles endogenous *vit-3* expression, and thus it is the primary vitellogenesis reporter used in this study.

**Figure 1.**
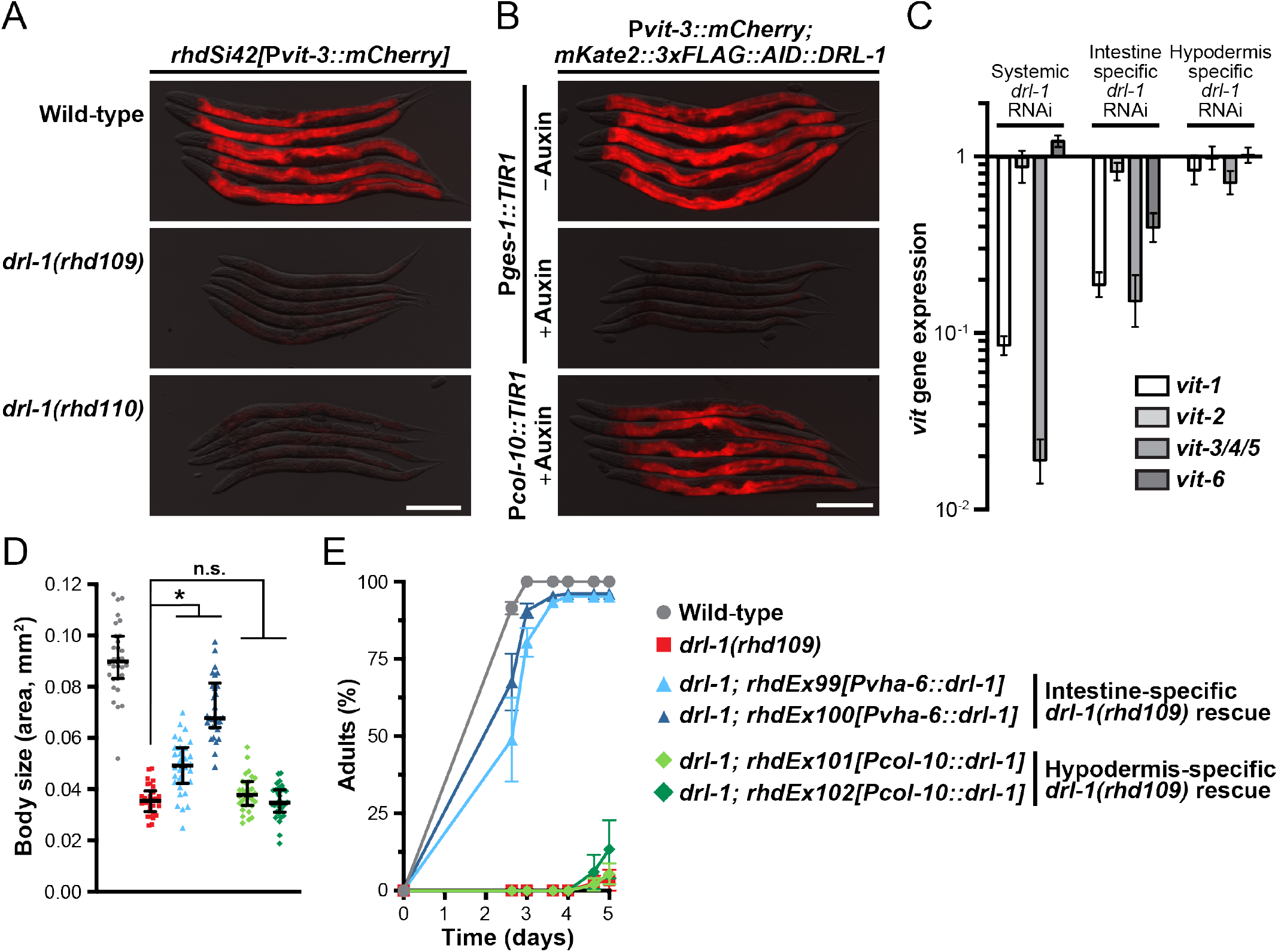
DRL-1 functions in the intestine to promote development, growth, and lipid reallocation. Representative overlaid DIC and mCherry fluorescence images of day 1 adult animals expressing a single-copy P*vit-3::mCherry* reporter (*vit-3* promoter fused to *mCherry*) in (A) wild-type or *drl-1* mutant animals or (B) *mKate2::3xFLAG::AID::drl-1* animals following intestinal (P*ges-1::TIR1*) or hypodermal (P*col-10::TIR1*) degradation using 4 mM auxin (scale bars, 200 μm). (C) RT-qPCR analysis of endogenous *vit* gene expression in day 1 adult animals after whole-body or tissue-specific knockdown of *drl-1* by RNAi. (D) Body size and (E) growth rate of wild-type, *drl-1(rhd109)*, or the indicated *drl-1(rhd109)* tissue-specific rescue strains (2 independent lines each). All strains contain *mgIs70* and the data are presented as (D) the median and interquartile range (*, *P*<0.0001, one-way ANOVA) or (E) the mean +/- SEM of three independent experiments.

We previously performed an RNAi screen for factors that are required for proper vitellogenin expression at the onset of adulthood and identified the *drl-1* gene as essential for robust P*vit-3::GFP* expression (Dowen et al. 2016). The *drl-1* gene encodes a protein kinase with highest similarity to the mammalian Mitogen-activated protein kinase kinase kinase 3 (MAP3K3/MEKK3) protein (Chamoli et al. 2014). To eliminate the possibility that *drl-1* RNAi produces off-target effects that impairs vitellogenesis, we generated several *drl-1* genetic mutants, which are all likely full loss-of-function alleles, using CRISPR/Cas9 gene editing and assessed *Pvit-3* reporter expression. Indeed, loss of *drl-1* resulted in a dramatic reduction in vitellogenin reporter expression (Fig. 1A, S1A-B). These findings are consistent with the observation that knockdown of *drl-1* reduces intestinal lipid stores (Chamoli et al. 2014), which could impair lipoprotein synthesis and assembly. Vitellogenin gene expression is regulated through non-cell-autonomous and cell-autonomous mechanisms via hypodermal, germline, and intestinal regulators (Perez and Lehner 2019), which prompted us to test where *drl-1* functions in the worm to regulate *vit* gene expression. After introduction of an auxin-inducible degron (AID) tag into the endogenous *drl-1* locus using CRISPR/Cas9 genome editing (Zhang et al. 2015; Ashley et al. 2021), we performed tissue-specific DRL-1 protein depletion in hypodermal or intestinal cells and assessed P*vit-3::mCherry* reporter expression. DRL-1 depletion in the intestine, but not the hypodermis, markedly impaired reporter expression (Fig. 1B), indicating that DRL-1 functions cell-autonomously to control vitellogenesis. Moreover, knockdown of *drl-1* in the intestine, but not in the hypodermis, using tissue-specific RNAi reduced the expression of the endogenous *vit* genes (Fig. 1C). Consistently, we were able to rescue vitellogenin reporter expression in *drl-1* mutant animals with a transgene that expresses *drl-1* under the control of an intestinal, but not a hypodermal, promoter (Fig. S1C), indicating intestinal *drl-1* expression is sufficient to restore vitellogenin expression.

Impaired lipid homeostasis can have profound effects on organismal growth and many mutants with vitellogenesis phenotypes also display gross defects in development (Perez and Lehner 2019). Thus, we inspected the *drl-1* mutants for growth rate and body size phenotypes, finding severe defects (Fig. S1D-E). Consistent with our previous observations, genetic rescue of the *drl-1* mutant with intestinal, but not the hypodermal, *drl-1* expression strongly suppressed these developmental phenotypes (Fig. 1D-E). Together, these data demonstrate an intestinal role for DRL-1 in the regulation of lipid allocation, growth rate, and body size.

### Intestinal DRL-1 and FLR-4 interact to form a protein kinase complex

The *flr-4* gene encodes a MAP Kinase that is required for metabolic homeostasis and proper aging, and importantly, mutants display a diet-specific extended lifespan that is similar to that of *drl-1* mutants (Chamoli et al. 2014; Verma et al. 2018), suggesting that these two MAP Kinases may act together to govern lipid homeostasis, growth, and development. To investigate this possibility, we first inspected the kinase domain of DRL-1 and FLR-4, as well as several other similar MAPKs, and discovered that while DRL-1 shares significant sequence similarity in the kinase domain it lacks several of the conserved amino acids that participate in ATP binding and catalysis (Fig. 2A). To address whether DRL-1 possesses kinase activity we used CRISPR/Cas9 to generate a DRL-1 P269S mutation, which is analogous to the FLR-4 P223S loss-of-function mutation in the activation loop of the kinase, and inspected P*vit-3::GFP* reporter expression. Similar to the *flr-4(P223S)* mutant, *drl-1(P269S)* animals display temperature-sensitive vitellogenesis defects (Fig. 2B, S2A). Additional mutation of E253 and G254 in the kinase domain, which are required for *in vitro* kinase activity (Chamoli et al. 2014), enhanced the temperature-sensitive vitellogenesis phenotype to similar levels as the null mutants (Fig. S2A). Not only is the kinase activity of DRL-1 and FLR-4 necessary for proper *vit* gene expression, but it is required to suppress the expression of the β-oxidation gene *ech-9* and the detoxification gene *ugt-18* (Fig. 2B) (Chamoli et al. 2014), indicating that DRL-1 and FLR-4 function broadly in regulation of intestinal metabolism.

**Figure 2.**
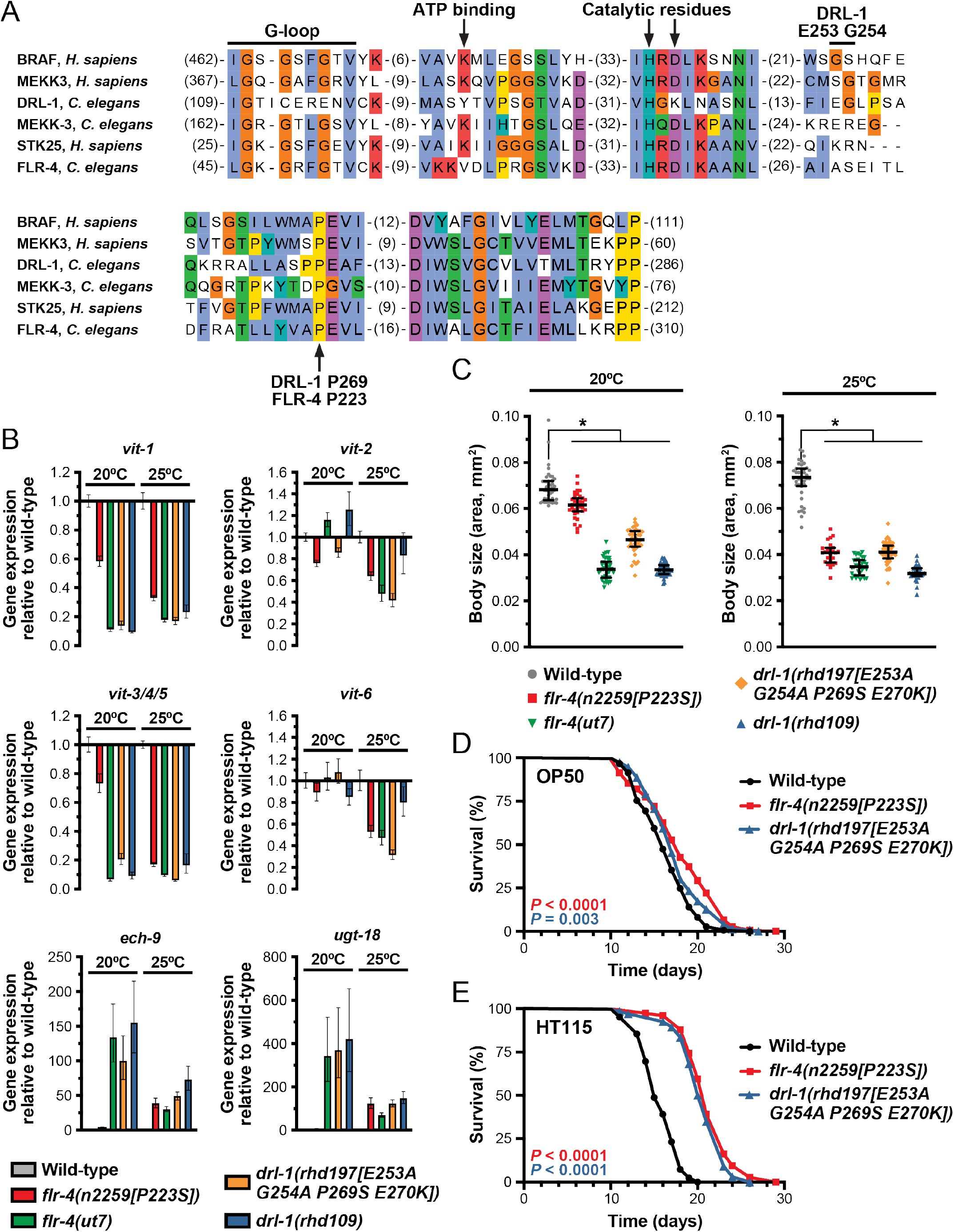
DRL-1 and FLR-4 kinase activity is essential for function. (A) An amino acid alignment (MUSCLE) of the kinase domain of the indicated MAP kinases. The key residues are indicated, including those mutated in the kinase dead mutants (DRL-1 E253, G254, P269; FLR-4 P223). (B) Expression of the indicated genes as measured by RT-qPCR in day 1 adult animals grown at 20 or 25**°**C. (C) Body size of the indicated strains at 20 or 25**°**C (*, *P*<0.0001, one-way ANOVA). (B, C) The *drl-1(rhd109)* and *flr-4(ut7)* alleles are presumed to be null mutations. (D, E) Longitudinal lifespan assays of wild-type and kinase dead mutants reared at 25**°**C with FUDR on *E. coli* (D) OP50 or (E) HT115. Log-rank test *P* values are reported.

Similar to the *drl-1* null mutants, *flr-4* mutants display severely reduced developmental rates and body size when reared on *E. coli* OP50, the standard laboratory diet (Fig. S2B-C) (Katsura et al. 1994; Verma et al. 2018). Furthermore, the kinase activity of DRL-1 and FLR-4 is required to maintain proper body size (Fig. 2C), suggesting that these kinases may function in a broader signaling cascade to balance metabolic needs during development and aging. Consistent with this hypothesis, knockdown of *flr-4* by RNAi extends lifespan when animals are reared on *E. coli* HT115, but not OP50 (Verma et al. 2018). To investigate whether DRL-1 kinase activity restricts lifespan, we performed longitudinal lifespan assays on the *flr-4(P223S)* and *drl-1(P269S)* mutants on different *E. coli* food sources. Indeed, DRL-1 and FLR-4 kinase dead mutants are markedly long-lived on *E. coli* HT115 (Fig. 2D-E). Together, these data demonstrate a role for DRL-1/FLR-4 kinase activity in maintaining overall organismal homeostasis.

These observations support the intriguing possibility that DRL-1 and FLR-4 act in the same tissue (*i*.*e*., the intestine), or even potentially in a protein complex, to coordinate developmental or nutritional programs. To investigate this hypothesis, we first sought to define where FLR-4 functions, focusing on the intestinal and neuronal tissues (Verma et al. 2018; Take-uchi et al. 2005). Using CRISPR/Cas9 to introduce an AID tag at the endogenous *flr-4* locus, we performed tissue-specific depletion experiments and assessed vitellogenin expression, body size, and growth rate. Depletion of FLR-4 in the intestine, but not in neurons, abrogated P*vit-3::mCherry* expression, severely reduced body size, and dramatically slowed growth rates (Fig. 3A-C, S3). Moreover, depletion of either DRL-1 or FLR-4 in the intestine is sufficient to confer unique responses to different *E. coli* food sources, including HT115-dependent effects on growth rate and lifespan (Fig. S4). Taken together, these data indicate that FLR-4 acts cell-autonomously to regulate intestinal homeostasis, possibly by directly interacting with DRL-1.

**Figure 3.**
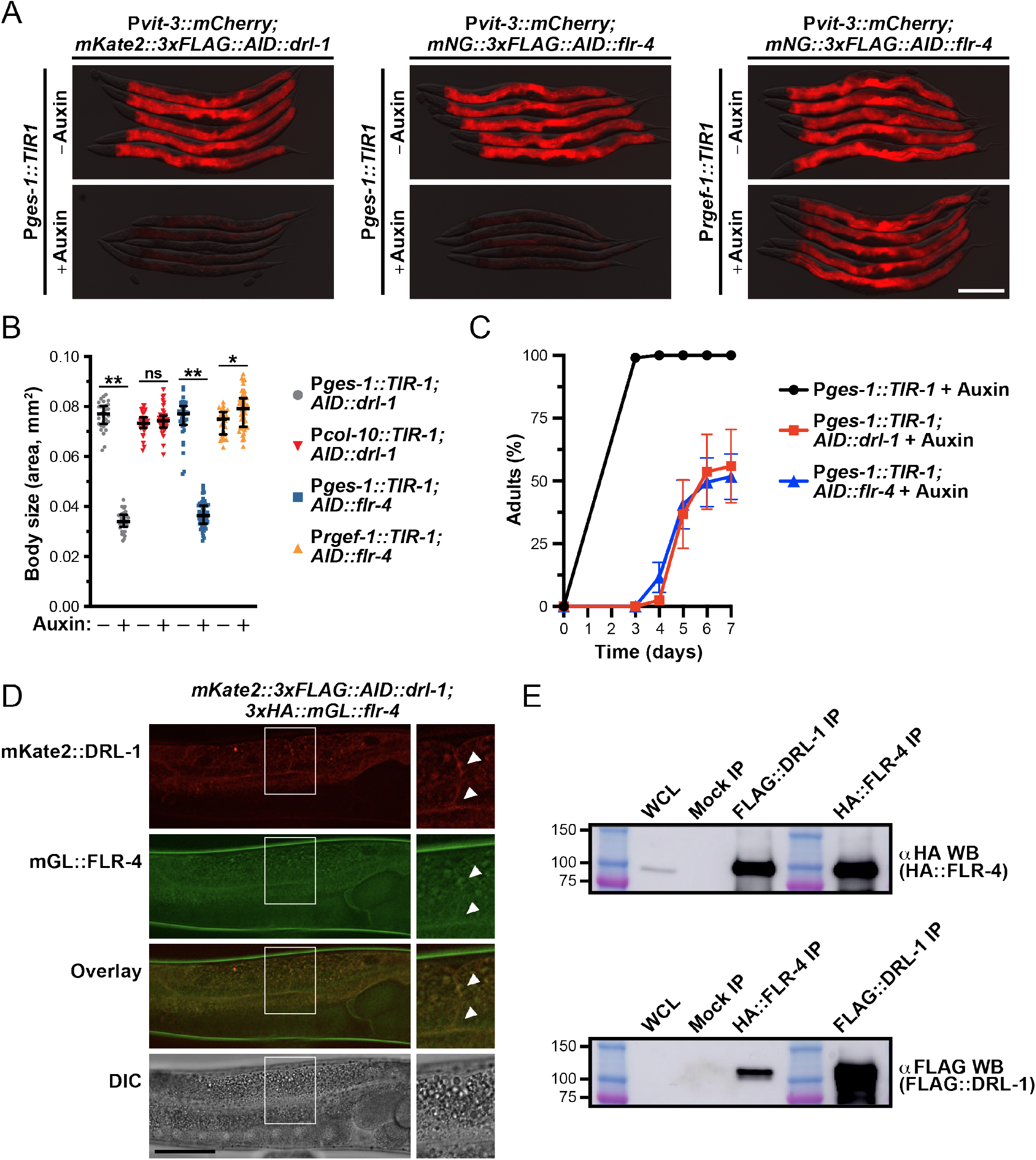
Intestinal DRL-1 and FLR-4 function in a complex to promote growth and lipoprotein production. (A-C) Phenotypic characterization of *mKate2::3xFLAG::AID::drl-1* or *mNG::3xFLAG::AID::flr-4* animals carrying the P*vit-3::mCherry* reporter after tissue-specific depletion with 4 mM auxin (P*ges-1::TIR1*, intestinal depletion; P*rgef-1::TIR1*, pan-neuronal depletion; P*col-10::TIR1*, hypodermal depletion). (A) Representative overlaid DIC and mCherry fluorescence images of day 1 adults with or without auxin treatment (scale bar, 200 μm), (B) body size of day 1 adults (median and interquartile range; ns, not significant, *, *P*=0.003, **, *P*<1×10^−45^, T-test), and (C) growth rate (mean +/-SEM) of animals after tissue-specific protein depletion. (D) Co-localization of mKate2::3xFLAG::DRL-1 and 3xHA::mGL::FLR-4 in intestinal cells of *glo-4(ok623)* animals (scale bar, 50 μm). Panels on the right show blowup images of the outlined regions and arrowheads point to areas of strong co-localization. (E) Co-immunoprecipitation of the indicated proteins after mock, anti-FLAG, or anti-HA immunoprecipitations followed by Western blotting. As a positive control, immunoprecipitations were probed with the same antibodies (far right lanes).

To test whether DRL-1 and FLR-4 form a protein complex, we used CRISPR/Cas9 gene editing to generate a strain that co-expresses HA::mGreenLantern::FLR-4 and mKate2::3xFLAG::DRL-1. Although these proteins are expressed at low levels, we detected the mGL::FLR-4 and mKate2::DRL-1 proteins at the intestinal plasma membrane (Fig. 3D), consistent with over-expression studies of FLR-4 and DRL-1 (Kobayashi et al. 2011; Wimberly and Choe 2022). Furthermore, reciprocal co-immunoprecipitation studies demonstrated that HA::FLR-4 and FLAG::DRL-1 interact (Fig. 3E), likely forming a larger protein kinase complex. These data explain why *drl-1* and *flr-4* mutants have similar phenotypes and provide a mechanistic basis for their role in maintaining intestinal homeostasis via MAPK signaling.

### The FLR-2 neuropeptide hormone antagonizes DRL-1 and FLR-4 signaling

DRL-1/FLR-4 signaling promotes organismal development, growth, and aging. Yet, it is possible that pro-growth signaling through this pathway may be dynamically tuned in response to adverse environmental or nutritional conditions to temper development. It is likely that this signaling would need to be balanced by other signaling events to maintain overall homeostasis. Thus, we reasoned that in the absence of *drl-1*/*flr-4* other pathways may actively restrict development. To identify components of this signaling axis, we performed a forward genetic screen to isolate mutations that suppress the growth and vitellogenesis defects associated with the *drl-1(rhd109)* mutant. We isolated over 100 mutants and initially pursued a small pilot set for further analysis. Following backcrossing and whole genome sequencing, we identified a putative null mutation in the *flr-2* gene as a likely *drl-1* suppressor mutation (Table S1).

The *flr-2* gene encodes a neuronally expressed secreted protein with highest similarity to the human glycoprotein hormone subunit α2 (GPA2) of thyrostimulin (Park et al. 2005; Oishi et al. 2009), as well as weaker similarity to the α subunits of other human glycoprotein hormones (*i*.*e*., FSH, LH, and TSH). The *flr-2(rhd117)* mutation identified in our screen, as well as a second allele *flr-2(ut5)*, strongly suppressed the defects in vitellogenin reporter expression observed in the *drl-1* and *flr-4* mutants (Fig. 4A, S5A). Furthermore, while loss of *drl-1* impaired the accumulation of neutral lipids as revealed by reduced Oil Red O staining (Chamoli et al. 2014), the *drl-1; flr-2* double mutants displayed wild-type fat levels (Fig. 4B, S5B). Consistent with these defects in vitellogenesis and lipid homeostasis, the *drl-1* mutant animals also had a dramatically smaller brood size; however, *drl-1; flr-2* double mutants produced nearly as many progeny as wild-type animals (Fig 4C). Together, these data suggest that FLR-2 opposes MAPK signaling to balance intestinal resources to maintain homeostasis.

**Figure 4.**
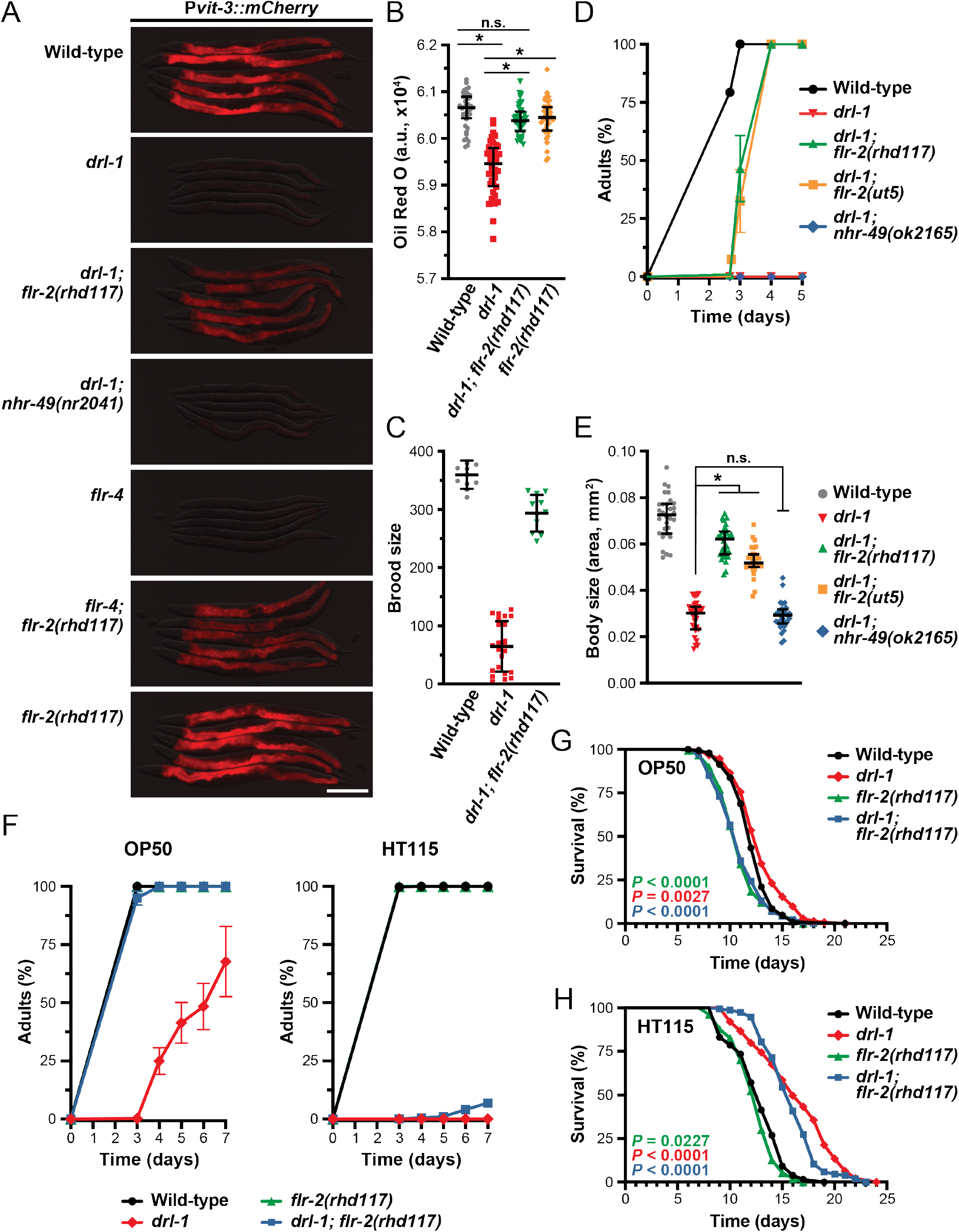
Loss of *flr-2* suppresses mutations in *drl-1* and *flr-4* when reared on specific food sources. (A) Representative fluorescence images of P*vit-3::mCherry* reporter expression (day 1 adults; scale bar, 200 μm), (B) whole animal Oil Red O staining (day 1 adults; median and interquartile range; ns, not significant, *, *P*<0.0001, one-way ANOVA), (C) brood size (mean +/- SD), (D) growth rate (mean +/- SEM), and (E) body size (day 1 adults; median and interquartile range; ns, not significant, *, *P*<0.0001, one-way ANOVA) for wild-type animals and the indicated mutants grown on *E. coli* OP50. (D, E) Strains contain the *mgIs70* transgene except for the *drl-1; nhr-49(ok2165)* strain. (F) The *flr-2(rhd117)* mutation robustly suppresses *drl-1* mutant growth defects when animals are reared on *E. coli* OP50 (left), but not *E. coli* HT115 (right). Longitudinal lifespan assays of wild-type animals and the indicated mutants reared at 25**°**C with FUDR on *E. coli* (G) OP50 or (H) HT115. Animals were L4s at day 0 and *P* values (log-rank test) are reported. (A-H) The *drl-1(rhd109)* and *flr-4(ut7)* alleles were used in these studies.

To assess whether FLR-2 impairs growth upon loss of *drl-1*, we performed growth measurements in the *drl-1* and *drl-1; flr-2* mutants. Indeed, both *flr-2* alleles suppress the slow growth rate and small body size phenotypes displayed by the *drl-1* mutant (Fig. 4D-E). Mutations in the nuclear hormone receptor *nhr-49* (orthologous to human HNF4 and PPARα), which has been previously shown to suppress *drl-1* RNAi phenotypes (Chamoli et al. 2014), failed to suppress the vitellogenesis, growth rate, and body size defects of the *drl-1(rhd109)* mutant (Fig. 4A, D-E). It is possible that experimental differences, such as the type of food or degree of *drl-1* inactivation (RNAi vs. mutant), account for these discrepancies.

Expression of the FLR-2 protein is limited to a small set of neurons in the head and tail (Oishi et al. 2009), suggesting that FLR-2 functions non-cell-autonomously to regulate intestinal homeostasis. Indeed, genetic rescue of *flr-2* under the control of a pan-neuronal promoter partially reverses the growth and body size phenotypes of the *drl-1; flr-2* double mutants (Fig. S5C-D), suggesting that expression of *flr-2* in neurons is sufficient to slow the growth of *drl-1* mutant animals. Together, these data demonstrate that the FLR-2 neurohormone is a potent inhibitor of growth, development, and reproduction in the absence of active DRL-1/FLR-4 signaling.

The lifespan and stress responses of *drl-1* and *flr-4* mutants are markedly different when animals are fed *E. coli* HT115 bacteria compared to OP50, likely due to differences in the nutritional value of the strains (Nair et al. 2022). Consistent with these observations, genetic mutation of *drl-1*, as well as intestinal depletion of AID::DRL-1, more severely impairs growth of animals reared on *E. coli* HT115 compared to OP50 (Fig. 4F, S4A). Although mutation of *flr-2* suppresses the growth defects of the *drl-1(rhd109)* mutant on OP50, it surprisingly fails to suppress *drl-1(rhd109)* growth on HT115 (Fig. 4F). Similarly, the *flr-2* mutation fails to suppress the longevity conferred by loss of *drl-1* when animals are grown on *E. coli* HT115 (Fig. 4G-H). Notably, loss of *flr-2*, which results in short-lived animals (Oishi et al. 2009), suppresses the modest lifespan increase displayed by *drl-1* mutant animals on OP50 (Fig. 4G). Our results demonstrate that while loss of *flr-2* strongly suppress the *drl-1* mutant phenotypes on OP50, it fails to yield similar results on HT115, suggesting that a HT115-specific nutritional input may be acting redundantly with FLR-2 to oppose DLR-1/FLR-4 signaling.

### FLR-2 acts via the G protein-coupled receptor FSHR-1 to stimulate PKA activity

The FLR-2 protein is a secreted hormone that likely acts by binding a cell surface receptor on a distal tissue, possibly the intestine, to stimulate a signaling cascade that slows animal development. We predicted that mutations in the FLR-2 receptor may also suppress the growth and vitellogenesis defects displayed by the *drl-1* mutant. Indeed, sequencing of additional *drl-1* suppressor mutations identified a putative null mutation in the *fshr-1* gene (Table S2), which encodes a G protein-coupled receptor with similarity to the family of glycoprotein hormone receptors that include the TSH, FSH, and LH receptors in humans (Cho et al. 2007). The *fshr-1(rhd118)* mutation, as well as the well-characterized *fshr-1(ok778)* allele, both suppressed the vitellogenesis defects conferred by loss of either *drl-1* or *flr-4* (Fig. 5A, S6A). Furthermore, the *fshr-1* mutations suppressed the slow growth and reduced body size of the *drl-1(rhd109)* mutant (Fig. 5B-C). The partial suppression of the body size can be attributed to the fact that *flr-2* and *fshr-1* single mutants are smaller than wild-type animals (Fig. S5D, S6B). We reasoned that if FSHR-1 is the intestinal receptor for FLR-2, then knockdown of *fshr-1* specifically in the intestine would suppress the *drl-1* mutation. Indeed, intestine-specific knockdown of *fshr-1* suppresses the vitellogenesis defects and small body size of *drl-1* mutant animals to similar levels as does systemic *fshr-1* knockdown (Fig. 5D-E). Together, these data suggest that intestinal FSHR-1 mediates the effects of FLR-2 to slow developmental rate of animals when DRL-1/FLR-4 signaling is reduced.

**Figure 5.**
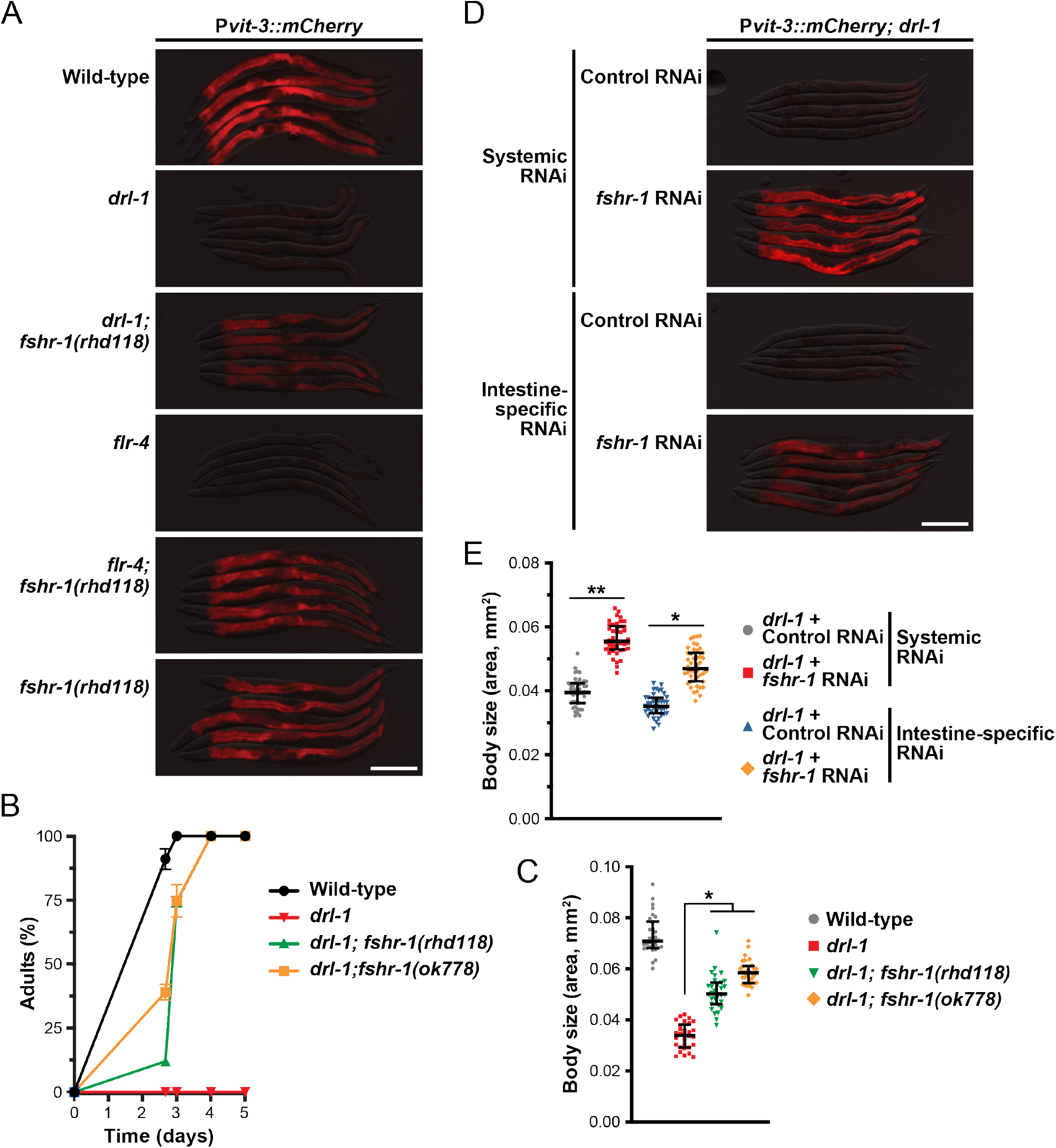
The intestinal GPCR FSHR-1 opposes DRL-1/FLR-4 signaling. (A) Representative overlaid DIC and mCherry fluorescence images (day 1 adults; scale bar, 200 μm), (B) growth rate (mean +/- SEM), and (C) body size (day 1 adults; median and interquartile range; *, *P*<0.0001, one-way ANOVA) for wild-type and mutant animals. (B, C) Strains contain the *mgIs70* transgene. (D) P*vit-3::mCherry* reporter expression (scale bar, 200 μm) and (E) body size (median and interquartile range; *, *P*<1×10^−19^, **, *P*<1×10^−30^, T-test) of day 1 adult *drl-1(rhd109)* animals subjected to systemic or intestine-specific RNAi.

The mammalian FSH receptor, like other glycoprotein hormone receptors, signals through heterotrimeric G proteins, primarily Gα_s_, to stimulate adenylate cyclase activity, cAMP production, and protein kinase A (PKA) activation (Dufau 1998; Casarini and Crépieux 2019). In *C. elegans*, gain-of-function mutations in *gsa-1* (Gα_s_) or *acy-1* (adenylate cyclase) suppresses the germline defects observed in the *fshr-1* mutant, providing genetic evidence that this pathway functions in the worm. Thus, we predicted that loss of *gsa-1*, the *acy-1-4* genes (adenylate cyclase), or *kin-1* (PKA) would also suppress the *drl-1* mutant phenotypes if FSHR-1 couples to this signaling pathway (Fig. 6A). RNAi knockdown of *gsa-1, acy-4*, or *kin-1* reactivated vitellogenin reporter expression to different degrees in the *drl-1* mutant background (Fig. 6B). Since FSHR-1 functions in the intestine to oppose DRL-1 signaling, we then performed intestine-specific RNAi against *gsa-1, acy-4*, and *kin-1* in the *drl-1* mutant and measured P*vit-3::mCherry* expression levels and body size (Fig. 6C-D). Indeed, inactivation of this canonical PKA activation pathway in the intestine suppressed the *drl-1* mutation to levels similar to *fshr-1* knockdown. The ligand for FSHR-1 has remained elusive despite its widespread roles in immunity, stress responses, and germline development (Cho et al. 2007; Powell et al. 2009; Kim and Sieburth 2020); however, our results are consistent with the possibility that FLR-2 is a ligand for FSHR-1 and induces cAMP signaling and PKA activation in the intestine.

**Figure 6.**
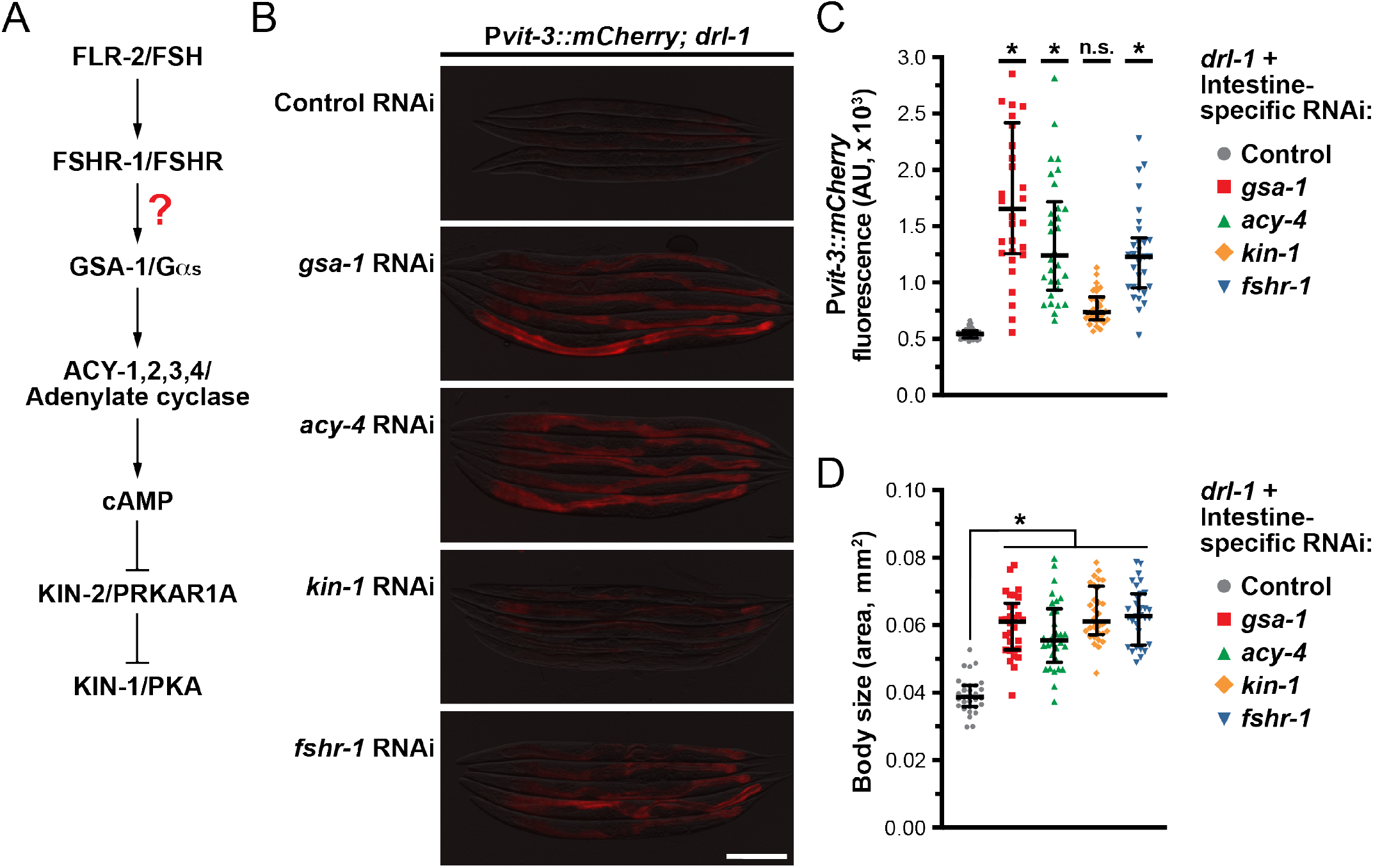
FLR-2 and FSHR-1 function through intestinal Gα_s_/cAMP signaling. (A) A diagram of how FLR-2/FSHR-1 may activate PKA via Gαs/cAMP signaling. (B) Representative overlaid DIC and mCherry fluorescence images of *drl-1(rhd109)* day 1 adults following systemic RNAi (scale bar, 200 μm). (C) Quantification of P*vit-3::mCherry* reporter expression and (D) body size of *drl-1(rhd109)* animals subjected to intestine-specific RNAi (day 1 adults; median and interquartile range; ns, not significant, *, *P*<0.0001, one-way ANOVA). (C, D) The intestine-specific RNAi was performed for one generation apart from *kin-1*, which was performed for two generations.

### FLR-2 and DLR-1/FLR-4 inversely regulate p38 signaling to tune development

It is possible that FLR-2/FSHR-1/PKA signaling functions in parallel to DRL-1/FLR-4 to differentially regulate a core developmental pathway. Intriguingly, mutations in components of the p38 MAPK pathway, including *tir-1* (orthologue of the human TIR domain protein SARM1), *nsy-1* (MAPKKK), *sek-1* (MAPKK), or *pmk-1* (MAPK), suppresses the increased lifespan of *flr-4* and *drl-1* mutants (Verma et al. 2018; Chamoli et al. 2020). The p38/PMK-1 pathway functions broadly in innate immunity (Kim et al. 2002; Aballay et al. 2003), response to oxidative stress (Inoue et al. 2005), development (Kim et al. 2016; Cheesman et al. 2016; Foster et al. 2020), and longevity (Wu et al. 2019). Moreover, loss of *flr-4* or *drl-1* results in hyperphosphorylation of PMK-1 (Verma et al. 2018; Chamoli et al. 2020), which may promote slower developmental rates (Cheesman et al. 2016). Consistent with these previous observations, our loss-of-function mutations in *drl-1* or *flr-4*, as well as intestinal depletion of AID::DRL-1 or AID::FLR-4 with auxin, increased the levels of active, phosphorylated PMK-1 (Fig. S7).

We hypothesized that hyperactivation of p38/PMK-1 may underlie the reduced developmental rates, small body sizes, and impaired vitellogenin production of the *drl-1* and *flr-4* mutants. Indeed, knockdown of the p38 pathway components *tir-1, nsy-1, sek-1*, or *pmk-1* by RNAi restored P*vit-3:mCherry* expression and increased the body size of *drl-1* mutant animals to varying degrees (Fig. S8A-B). Moreover, intestine-specific knockdown of *pmk-1* partially suppressed the vitellogenesis defects, small body size, and slow growth of the *drl-1* mutant (Fig. 7A-C). Intestinal knockdown of *pmk-1* also partially suppressed the body size defects of animals depleted of intestinal FLR-4 (Fig. S8C). Importantly, loss of *flr-2* was significantly more effective in suppressing DRL-1 depletion than a null mutation in *pmk-1* (Fig. 7D), suggesting that FLR-2/FSHR-1 may function through multiple parallel pathways to modulate growth and lipid homeostasis.

**Figure 7.**
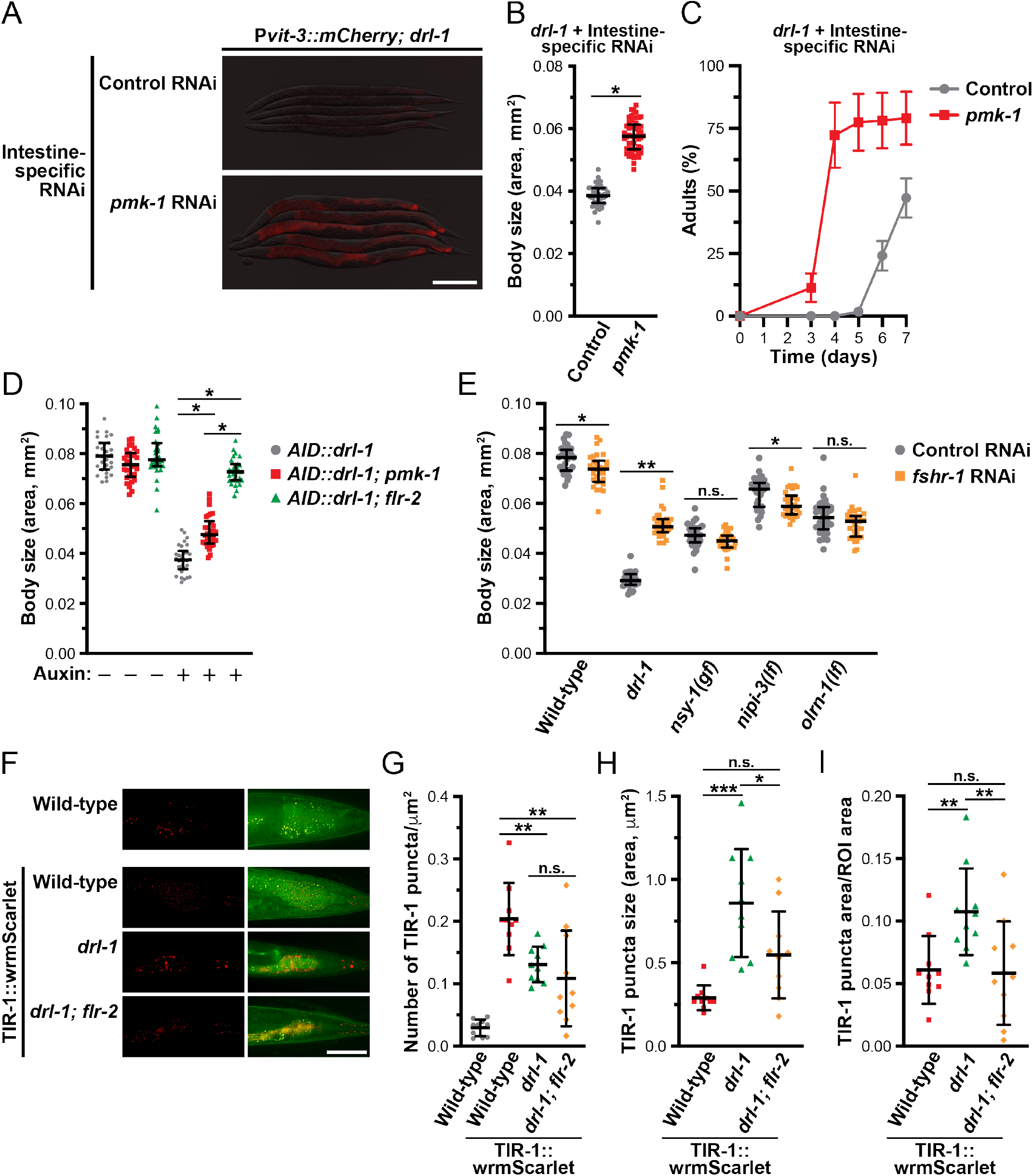
DRL-1 and FLR-2/FSHR-1 signaling balance p38/PMK-1 activity to promote growth and development. (A) Representative overlaid DIC and mCherry fluorescence images of day 1 adults (scale bar, 200 μm), (B) body size (day 1 adults; median and interquartile range; *, *P*=3×10^−36^, T-test), and (C) growth rate (mean +/- SEM) of *drl-1(rhd109)* animals subjected to intestine-specific RNAi. (D) Body size of *AID::drl-1* day 1 adult animals following intestinal depletion of DRL-1 with 4 mM auxin (median and interquartile range; *, *P*<0.0001, one-way ANOVA). (E) Body size of wild-type and the indicated mutants after control or *fshr-1* RNAi (median and interquartile range; *, *P*<0.01, **, *P*<0.0001, T-test). These mutations have been previously shown to activate p38/PMK-1 signaling. (F) Representative images of TIR-1::wrmScarlet localization in posterior intestinal cells (scale bar, 50 μm). The red channel shows the TIR-1 puncta (left panels), the green/red overlay shows yellow puncta that result from intestinal autofluorescence (right panels), and wild-type animals lacking TIR-1::wrmScarlet are included as a negative control (top panels). Quantification of the (G) total number, (H) absolute size, and (I) relative size of TIR-1 puncta in day 1 adult wild-type and *drl-1(rhd109)* mutant animals (mean +/- SD; ns, not significant, *, *P*<0.05, **, *P*<0.01, ***, *P*<0.0001, one-way ANOVA).

Our results suggest that DRL-1/FLR-4 and FLR-2/FSHR-1 exert opposing inhibitory and stimulatory effects, respectively, on the p38 signaling pathway to balance intestinal resources between developmental programs and stress responses. Consistent with this model, a gain-of-function mutation in Gα_s_, *gsa-1(ce81)*, or a loss-of-function mutation in the inhibitory regulator of PKA, *kin-2(ce179)*, reduced body size (Schade et al. 2005); however this phenotype is partially suppressed by *pmk-1* RNAi (Fig. S8D), suggesting that PKA likely functions upstream of PMK-1.

Although our genetic data argue that FLR-2/FSHR-1/PKA activates the p38 pathway, the site of this regulation is unknown. The MAP3K gain-of-function (gf) mutation, *nsy-1(ums8)*, hyperactivates downstream p38 signaling, resulting in developmental delay and small body size (Cheesman et al. 2016). Using this mutant, we tested whether *fshr-1* functions genetically upstream of *nsy-1*. While *fshr-1* RNAi strongly suppresses the *drl-1(rhd109)* mutation, it fails to suppress the small body size of the *nsy-1* gf mutant (Fig. 7E), indicating that FSHR-1 likely functions upstream of NSY-1 to regulate development. Moreover, other genetic perturbations that induce p38 hyperactivation include mutations in the neuronal developmental regulator *olrn-1* or the intestinal pseudokinase *nipi-3* (Kim et al. 2016; Foster et al. 2020; Wu et al. 2021); however, knockdown of *fshr-1* failed to suppress the body size defects caused by the *olrn-1* or *nipi-3* mutations, indicating that these pathways function independently of *fshr-1* (Fig. 7E).

Given that FSHR-1 functions upstream of NSY-1, we investigated whether the activity of TIR-1, the *C. elegans* orthologue of SARM1 (sterile alpha and TIR motif-containing 1) and upstream regulator of the p38 pathway (Liberati et al. 2004; Couillault et al. 2004), is modified by loss of *drl-1*. TIR-1 activation is triggered by a stress-induced phase transition that promotes protein oligomerization, NAD^+^ glycohydrolase activity, and stimulation of the downstream NSY-1/SEK-1/PMK-1 pathway (Loring et al. 2021; Peterson et al. 2022). Using a strain expressing TIR-1::wrmScarlet (Peterson et al. 2022), we tested whether loss of *drl-1* could enhance TIR-1 phase transition, which is visible as intestinal puncta that are distinct from the autofluorescent gut granules. While mutation of *drl-1* stimulated a reduction in the number of TIR-1::wrmScarlet puncta, the size of the puncta was markedly increased, suggesting that loss of *drl-1* induces TIR-1::wrmScarlet phase transition (Fig. 7F-I). This *drl-1*-induced phase transition of TIR-1 was suppressed by loss of *flr-2*. Together, these data suggest that DRL-1/FLR-4 and FLR-2/FSHR-1/PKA exert opposing effects on TIR-1 phase separation to govern the activity of p38 signaling and animal development.

While the transcription factors that act downstream of p38 signaling to regulate stress responses have been well studied in *C. elegans* (Pukkila-Worley and Ausubel 2012), those that function in p38-regulated development are not well understood. Thus, we performed a small-scale RNAi screen to identify transcriptional regulators that function downstream of *drl-1* to regulate development. Using the P*vit-3::mCherry* reporter, we first assessed whether knockdown of each transcription factor could suppress the vitellogenesis defects of the *drl-1* mutant. Interestingly, knockdown of *pha-4*, a FoxA transcription factor, partially suppressed the vitellogenesis and body size defects conferred by the *drl-1* mutation (Fig. 8A-B). PHA-4 functions broadly in *C. elegans* development, but also has a distinct role in promoting longevity in response to dietary restriction (Panowski et al. 2007). Furthermore, *pha-4* is required for the lifespan increase that results from loss of *drl-1* or *flr-4* (Chamoli et al. 2014; Verma et al. 2018). Thus, we reasoned that DRL-1/FLR-4 may impact the localization of PHA-4, which we assessed using a strain that expresses an endogenously-tagged PHA-4::GFP to avoid potential artifacts of PHA-4 over-expression. Indeed, depletion of DRL-1 stimulated the accumulation of PHA-4::GFP protein in the nucleus of intestinal cells (Fig. 8C-D). Moreover, the PHA-4 nuclear accumulation was dependent on *pmk-1* regardless of the food source, suggesting that DRL-1, and downstream p38/PMK-1 signaling, may regulate the nuclear translocation of PHA-4/FOXA (Fig. 8C-D, S9). Dynamic nuclear translocation of PHA-4 has not yet been observed in *C. elegans* (Panowski et al. 2007; Sheaffer et al. 2008); however, these previous studies used PHA-4 over-expression transgenes while our experiments employed a CRISPR/Cas9-based GFP knock-in at the *pha-4* locus, which tags all isoforms. Notably, the human FOXA proteins show different abilities to dynamically shuttle between the nucleus and cytoplasm (Wolfrum et al. 2003). Together, our data demonstrate that DRL-1 signaling governs TIR-1 phase transition to module p38 activity and PHA-4 localization in the intestine to tune the developmental rate of *C. elegans*.

**Figure 8.**
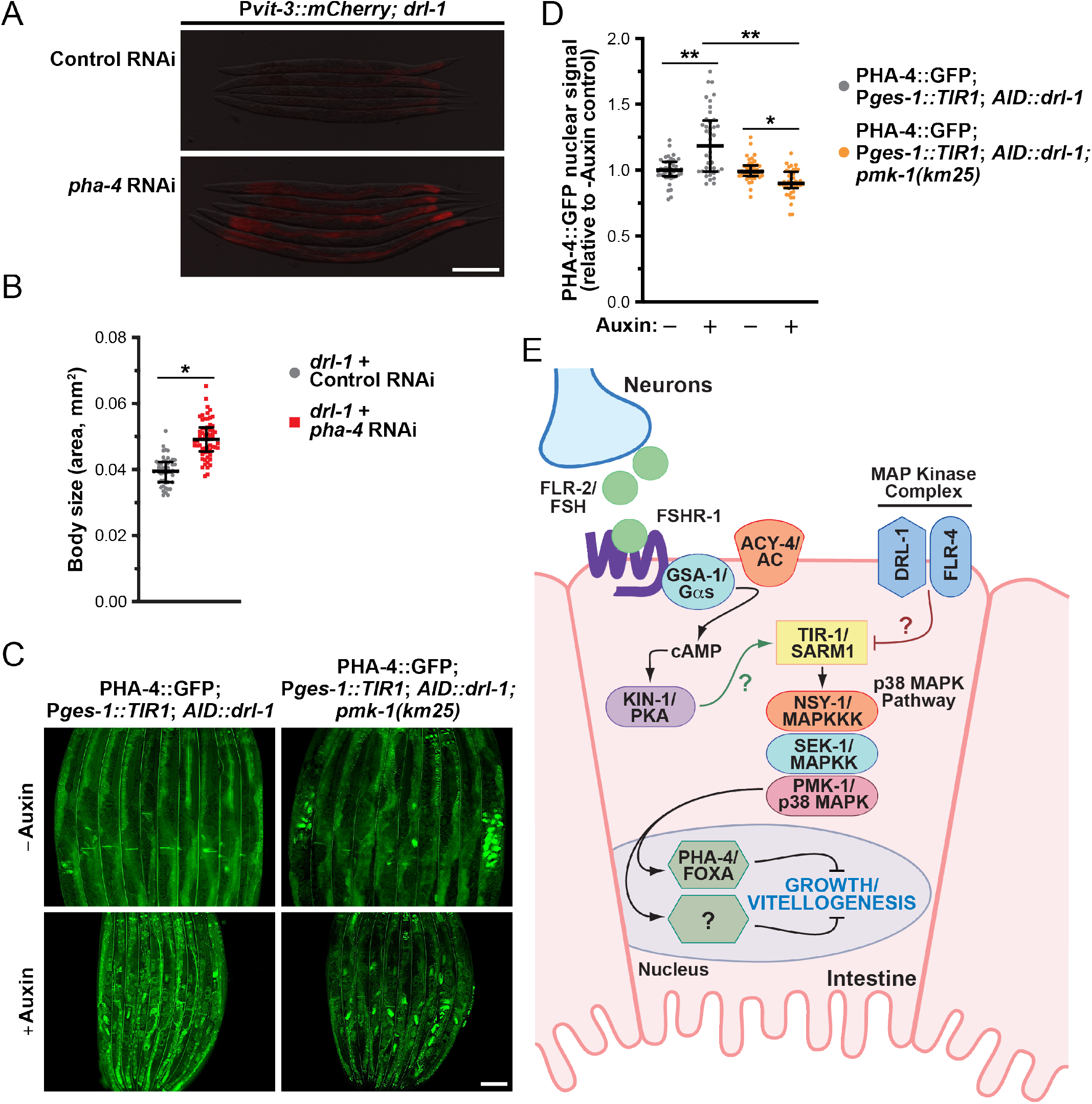
PHA-4/FOXA suppresses growth upon loss of *drl-1*. (A) Representative fluorescence images of P*vit-3::mCherry* reporter expression (F1 animals; scale bar, 200 μm) and (B) body size (P0 animals; median and interquartile range; *, *P*=7×10^−17^, T-test) of *drl-1(rhd109)* day 1 adults after knock-down of *pha-4* by RNAi. (B) This control RNAi data is also presented in Figure 5E. (C) GFP fluorescence images (scale bar, 100 μm) and (D) quantification (median and interquartile range; *, *P*<0.05, **, *P*<0.0001, one-way ANOVA) of PHA-4::GFP nuclear localization after intestine-specific depletion of AID::DRL-1 using 4 mM auxin in wild-type or *pmk-1(km25)* animals reared on *E. coli* OP50. (E) A model illustrating the opposing effects of DRL-1/FLR-4 and FLR-2/FSHR-1/PKA on p38-mediated growth.

## DISCUSSION

Here, we demonstrate that the DRL-1 and FLR-4 MAP kinases play crucial roles in managing organismal development, growth, and lipid homeostasis. The action of these pro-growth MAPKs, which form a protein complex on the plasma membrane of intestinal cells, is opposed by the neurohormone FLR-2, the *C. elegans* orthologue of the follicle stimulating hormone, and its putative intestinal G protein-coupled receptor FSHR-1 and downstream Gα_s_/cAMP/PKA signaling (Fig. 8E). In turn, MAPK and FSH signaling balance the activity of p38/PMK-1 by modulating the phase transition of TIR-1/SARM1, thereby tuning developmental rate and growth. At least some of the intestinal response to altered p38 activity is conferred at the level of transcription, as loss of *pha-4* partially rescues the developmental defects displayed by the *drl-1* mutant. Our study establishes a new mechanism by which the intestinal p38 signaling pathway integrates tissue-specific signals to govern organismal growth and development.

Our work suggests that loss of either *drl-1* or *flr-4* results in widespread misuse of energy resources, causing slow growth, small body size, and loss of fat storage. It is likely that impaired vitellogenin production is a consequence of this dietary restriction-like metabolic state, as other models of dietary restriction also display reduced vitellogenin levels (Seah et al. 2016). Notably, mutations that abrogate the kinase activity of DRL-1 or FLR-4 result in strong loss-of-function phenotypes, suggesting that both proteins may function in canonical MAPK kinase signaling in *C. elegans*. While the kinase domain of DLR-1 shares highest similarity to the mammalian MEKK3 protein (Chamoli et al. 2014; Wimberly and Choe 2022), the FLR-4 kinase domain is most similar to the catalytic domains of the GCKIII kinases (*i*.*e*., STK24/MST3, STK25/SOK1, and STK26/MST4) (Take-uchi et al. 2005). Interestingly, knockdown of STK25 increases β-oxidation and impairs lipid accumulation in human hepatocytes (Amrutkar et al. 2016), which is consistent with *flr-4* loss-of-function phenotypes.

Here, we show that DRL-1 and FLR-4 form a protein complex. While MEKK3 and GCKIII kinases are not known to directly interact, both have been shown to interact with the CCM (cerebral cavernous malformation) adaptor proteins (Faurobert and Albiges-Rizo 2010). Specifically, CCM2/OSM scaffolds MEKK3 at membranes where it activates p38 during osmotic stress (Uhlik et al. 2003; Zawistowski et al. 2005). In this cellular context MEKK3 promotes p38 activity, however in different cell types, CCM2 and CCM3 has been shown to have no impact on p38 activity or to negatively regulate p38 (Whitehead et al. 2009; Zhu et al. 2010), suggesting that cell type-specific scaffolds may be crucial to defining how the MEKK3 or the GCKIII kinases impact p38 activity. In *C. elegans*, the CCM orthologues KRI-1, CCM-2, and CCM-3 have poorly defined roles in intestinal metabolism and additional studies will be crucial to establishing whether DRL-1 and FLR-4 function in complex with these adaptor proteins.

While previous studies have established a genetic interaction between *drl-1*/*flr-4* and the p38 pathway (Verma et al. 2018; Chamoli et al. 2020), the molecular mechanism of this crosstalk has been unclear. We demonstrate a surprising mode of regulation whereby phase transition of TIR-1/SARM1 is repressed by DRL-1, and likely FLR-4 as well, to modulate downstream p38 signaling. Conversely, follicle stimulating hormone signaling promotes TIR-1 phase transition, likely through a FLR-2/FSHR-1/PKA signaling axis. Regulation of TIR-1/SARM1 phase transition via phosphorylation has not been demonstrated; however in human cells, phosphorylation of SARM1 by JNK promotes the intrinsic NAD^+^ hydrolase activity of the TIR domain (Murata et al. 2018). It’s possible that PKA and DRL-1/FLR-4 may individually phosphorylate TIR-1 to exert opposite effects on phase transition; or alternatively, DRL-1/FLR-4 may regulate PKA activity, which in turn phosphorylates TIR-1. Intriguingly, in cardiomyocytes STK25 inhibits PKA via phosphorylation of the regulatory subunit PRKAR1A (Zhang et al. 2022), suggesting that FLR-4 may function in a similar manner to regulate PKA in *C. elegans*.

While direct regulation of TIR-1 via phosphorylation is plausible, it is also possible that DRL-1/FLR-4 and FSH signaling act indirectly to control p38/PMK-1 signaling. A recent study demonstrated that cholesterol deficiency or loss of the NHR-8 nuclear hormone receptor, which is required for maintaining cholesterol homeostasis in *C. elegans*, stimulates p38 activity by promoting TIR-1 phase transition and NAD^+^ hydrolase activity (Magner et al. 2013; Peterson et al. 2022). Thus, loss of DRL-1 or FLR-4 could induce similar conditions of cholesterol mishandling, which would be consistent with the impaired intestinal lipid homeostasis, the altered cytoprotective responses, and heightened detoxification response that is observed in *drl-1* mutant animals (Chamoli et al. 2014; Wimberly and Choe 2022). Notably, loss of *drl-1* upregulates numerous cytochrome P450 and UDP-glucuronosyltransferase detoxification genes, which could deplete cholesterol stores by increasing flux through sterol modification and catabolic pathways (Chamoli et al. 2014; Larigot et al. 2022).

We demonstrate that the opposing action of FSH signaling and DRL-1/FLR-4 on p38/PMK-1 signaling governs the nuclear localization of PHA-4/FOXA in the intestine. To our knowledge, this is the first demonstration that PHA-4/FOXA nuclear localization is controlled by p38 signaling. The mammalian FOXA transcription factors are crucial regulators of early development and post-natal metabolic homeostasis (Carlsson and Mahlapuu 2002). Similarly, in *C. elegans* PHA-4 specifies pharyngeal cell fates and is required for the development of the foregut (Mango et al. 1994; Kalb et al. 1998; Horner et al. 1998), as well as post-embryonic regulation of metabolism and aging (Panowski et al. 2007; Sheaffer et al. 2008; Wu et al. 2018). Given the well-defined role of PHA-4 in promoting development, it is surprising that PHA-4 restricts, either directly or indirectly, vitellogenin production and body size upon loss of intestinal DRL-1. It is possible that phosphorylation by PMK-1 not only directs the nuclear localization of PHA-4 but also its preference for transcriptional targets. Moreover, it will be crucial to define the tissue of action and the developmental role of PHA-4 during dietary restriction (DR), as *pha-4* is required for DR-induced longevity (Panowski et al. 2007).

We found that *flr-2* mutations are stronger suppressors of *drl-1* mutant phenotypes than the *pmk-1* null mutation, suggesting that FSH signaling controls a p38-independent pathway. Consistently, intestinal FSHR-1 acts in parallel to the p38 and insulin signaling pathways to support the innate immune response (Powell et al. 2009). Moreover, the developmental defects, metabolic reprogramming, and misexpression of several cytoprotective genes induced by loss of *drl-1* are not strictly dependent on p38 signaling (Chamoli et al. 2020; Wimberly and Choe 2022). Thus, it is likely that the FLR-2/FSHR-1/PKA signaling pathway also promotes metabolic and developmental defects in *drl-1* mutants through a p38-independent mechanism. Notably, PKA promotes lipid mobilization in response to fasting and cold stress, likely by phosphorylating and stabilizing the ATGL-1 lipase on the surface of intestinal lipid droplets (Pagnon et al. 2012; Lee et al. 2014; Liu et al. 2017). Similar to the fasting response, knockdown of *drl-1* stimulates β-oxidation and upregulation of lipid catabolism genes (*e*.*g*., *cpt-3*/carnitine palmitoyl transferase), which in turn likely increases energy production through mitochondrial oxidative phosphorylation (Chamoli et al. 2014; Lee et al. 2014). Thus, it is possible that the p38-independent metabolic defects displayed by the *drl-1* mutant are a result of PKA-dependent lipid mishandling of intestinal lipid droplets. In the future, it will be crucial to define the metabolic role of FSH signaling in mammalian non-reproductive tissues, including the liver and adipose tissue, as well as assess whether MAPK signaling impinges on these pathways to integrate metabolic and developmental programs.

The DRL-1 and FLR-4 MAPKs are crucial to maintaining cellular homeostasis and balancing pro-growth programs against energy utilization. We propose that FSH and DRL-1/FLR-4 are likely to be dynamically regulated by nutritional inputs or environmental stimuli. Conceivably, the metabolic reprogramming and lifespan extension triggered by dietary restriction, or the elevated lipid utilization in response to short-term fasting, could engage neuronal FSH signaling to promote breakdown of intestinal lipids. Intriguingly, an *E. coli* HT115 diet further impairs the developmental rate of the *drl-1* mutant, and loss of *flr-2* fails to suppress this defect, suggesting that additional nutritional or sensing pathways may function in parallel with FSH signaling to restrict development (Nair et al. 2022). This work establishes the framework for identifying these unknown regulators, which will be vital to gaining a holistic view of the homeostatic mechanisms that promote development and reproductive fitness.

## MATERIALS AND METHODS

### C. elegans strains

*C. elegans* strains were cultured on NGM media seeded with *E. coli* OP50 or HT115(DE3) (Brenner 1974). Animals were reared at 20**°**C unless specified otherwise. For auxin-inducible degradation experiments, embryos were transferred to plates containing 4 mM Naphthaleneacetic Acid (K-NAA, PhytoTech) and grown to adulthood. All stains used in this study are listed in Supplemental Table S3.

### Generation and imaging of transgenic animals

Strains carrying the high-copy *mgIs70[*P*vit-3::GFP]* transgene or the single-copy *rhdSi42[*P*vit-3::mCherry]* transgene have been previously described (Dowen et al. 2016). The *drl-1* rescue constructs were generated by fusing the *col-10* promoter (chromosome V: 9,166,416–9,165,291; WS284) or the *vha-6* promoter (chromosome II: 11,439,355–11,438,422; WS284) to the *drl-1* cDNA (1,770bp of coding sequence with 141bp of 3’UTR) via Gibson assembly to generate plasmids pRD141 and pRD142, respectively (Gibson et al. 2009; Dowen et al. 2016). The resulting plasmids were microinjected into *drl-1(rhd109); mgIs70[Pvit-3::GFP]* animals at 20 ng/μl, along with 2.5 ng/μl pCFJ90(P*myo-2::mCherry*) and 77.5 ng/μl of 2-Log DNA ladder (New England BioLabs), to generate two independent strains expressing *Ex[*P*col-10::mCherry::his-58::SL2::drl-1 cDNA]* (DLS515, DLS516) and *Ex[*P*vha-6::mCherry::his-58::SL2::drl-1 cDNA]* (DLS513, DLS514). For the pan-neuronal *flr-2* rescue transgene, the *sng-1* promoter (chromosome X: 7,325,641–7,327,607; WS284) was fused to the *flr-2* gene (including the 3’ UTR) via Gibson assembly to generate the pRD157 plasmid. This plasmid, which is derived from the MosSCI-compatible pCFJ151 plasmid, was microinjected into EG6699 to generate the single-copy integrant *rhdSi46[*P*sng-1::mCherry::his-58::SL2::flr-2]* as previously described (Frøkjær-Jensen et al. 2008). All strains carrying *mgIs70* or *rhdSi42* were imaged on a Nikon SMZ-18 Stereo microscope equipped with a DS-Qi2 monochrome camera.

### CRISPR/Cas9 gene editing

Generation of the *drl-1* deletion alleles (*rhd109* and *rhd110*) were generated using the *pha-1* co-conversion approach, as previous described (Ward 2015). All additional edits were performed by microinjection of Cas9::crRNA:tracrRNA complexes (Integrated DNA Technologies) into the germlines of *C. elegans* animals as previously described (Ghanta and Mello 2020). Large dsDNA donor molecules with ∼40 bp homology arms on each end were prepared by PCR using Q5 DNA Polymerase (New England BioLabs) and purified using HighPrep PCR Clean-up beads (MagBio) per the manufacturers’ instructions. The DNA repair templates were melted and reannealed prior to microinjection (Ghanta and Mello 2020). The PCR templates used to generate the dsDNA donor molecules were pRD156, pBluescript II(*linker::mKate2::TEV::linker::3xFLAG::AID*); pRD160, pBluescript II(*AviTag::linker::2xTEV::linker::3xHA::linker::mNeonGreen::TEV::linker::3xFLAG::AID*); and pRD174, pBluescript II(*AviTag::linker::2xTEV::linker::3xHA::linker::mGreenLantern::TEV::linker::3xFLAG::AID*). To generate missense mutations using CRISPR/Cas9 gene editing, single-stranded oligodeoxynucleotides were used as donor molecules (Ghanta and Mello 2020). All CRISPR crRNA guide sequences are listed in Supplemental Table S4.

### RNAi Experiments

*E. coli* HT115(DE3) strains carrying the control L4440 plasmid or individual RNAi plasmids for gene expression knockdown were grown for ∼16 hrs in Luria–Bertani medium containing ampicillin (50 μg/ml), concentrated by 20-30x via centrifugation, and seeded on NGM plates containing 5 mM isopropyl-β-D-thiogalactoside (IPTG) and 50 μg/ml ampicillin. Plates seeded with RNAi bacteria were maintained at room temperature overnight to induce expression of the dsRNAs. All RNAi clones were selected from the Ahringer or Ahringer Supplemental RNAi library and confirmed by Sanger sequencing, with the exception of *nsy-1*, which has been described previously (Cheesman et al. 2016), and *sek-1* and *pmk-1*, which were generated by cloning cDNA fragments into the pL4440 vector using standard techniques. Synchronized *C. elegans* L1 larvae were generated by bleaching gravid animals to liberate embryos followed by overnight incubation in M9 buffer. Synchronized L1s were dropped on RNAi plates, grown at 20°C until they were day one adults (72-120 hrs), and processed for imaging.

### Growth, Lifespan, and Brood Size Assays

Animals were grown on their respective *E. coli* food sources (OP50 or HT115) for at least two generations without starvation prior to being assayed growth, lifespan, and brood size. For developmental growth rate assays, 100-200 eggs were picked from plates with freshly laid embryos (16 hrs or less) to new plates. Animals were scored every 24 hours for 7 days for the presence of gravid adults, which were promptly removed and recorded as having reached adulthood. For RNAi-treated animals, synchronized L1s were dropped on RNAi plates grown for up to 7 days and similarly scored. For developmental experiments employing auxin-inducible degradation of AID::DRL-1 or AID::FLR-4, freshly laid embryos were picked to plates containing 4 mM Naphthaleneacetic Acid and worm growth was scored as described above.

For body size measurements, L4 animals were picked to fresh plates and imaged 24 hrs later on a Nikon SMZ-18 Stereo microscope equipped with a DS-Qi2 monochrome camera. Using the Fiji software (Schindelin et al. 2012), animals were outlined by hand and the number of pixels were measured, which was converted to a square micron value based on the known imaging settings. Body size data in mm^2^ are presented as the median with the interquartile range. A one-way ANOVA with a Bonferroni correction for multiple testing was performed to determine whether samples were significantly different.

Lifespan assays were performed at 25°C in the presence of 50 µM FUDR unless otherwise noted. Briefly, well-fed L4 animals were picked to FUDR-containing plates and animals (30 animals/plate, 120-150 total animals) were maintained on uncontaminated plates and scored daily for survival as previously described (Dowen 2019). A log-rank test was applied to determine whether survival curves were significantly different.

For brood sizes experiments, L4 animals (N2, n=11; *drl-1(rhd109)*, n=23; *drl-1(rhd109); flr-2(rhd117)*, n=11) were picked to individual plates and transferred daily throughout the reproductive period. Progeny were counted as L3 or L4 animals and data are reported as the mean +/-the standard deviation.

### Oil Red O staining

Approximately 75 L4 animals were picked to new plates, harvested 24 hours later as day 1 adults in S buffer, washed, fixed with 60% isopropanol, and stained for 7 hours with 0.3% Oil Red O as previously described (Dowen 2019). Animals were mounted on 2% agarose pads and imaged with a Nikon SMZ-18 Stereo microscope equipped with a DS-Qi2 monochrome camera for intensity analyses or a Nikon Ti2 widefield microscope equipped with a DS-Fi3 for representative color images. Quantification of Oil Red O staining was performed in ImageJ by manually outlining worms and determining the mean gray value in the worm area. Each value was subtracted from 65,536, the maximum gray value for 16-bit images. The data were plotted using Prism 9 and a one-way ANOVA with a Bonferroni correction for multiple testing was performed to calculate *P* values.

### Quantitative PCR

L1 animals were synchronized by bleaching and grown to the first day of adulthood prior to being harvested in M9 buffer, washed, and flash frozen. The total RNA was isolated using Trizol (Thermo Fisher), followed by chloroform extraction and precipitation with isopropanol. DNase treatment, followed by cDNA synthesis with oligo(dT) priming, was performed using the SuperScript IV VILO Master Mix with ezDNase Kit according to the manufacturer’s instructions (Thermo Fisher). Quantitative PCR was performed exactly as previously described (Dowen 2019). All primer sequences are listed in Supplemental Table S5. Expression data are presented as the mean fold change relative to a control (N2 for mutant analyses, L4440 for RNAi analyses) with the Standard Error of the Mean (SEM) reported for three independent experiments.

### Western Blot Analyses

Animals were synchronized by bleaching and the resulting L1s were grown to day one adults. The animals were harvested in M9 buffer, washed three times, and the worm pellets were snap frozen in liquid nitrogen. The pellets were resuspended in an equal volume of 2x RIPA buffer (Cell Signaling Technology) containing a 2x Halt Protease and Phosphatase Inhibitor Cocktail (Thermo Fisher) and homogenized for approximately 15 seconds with disposable pellet pestle (Thermo Fisher) mounted in a cordless drill (Dewalt). Samples (∼200 µL) were then subjected to sonication (30 sec on/off cycles, 10 cycles) using a Bioruptor Pico sonication instrument (Diagenode). Whole cell lysates were cleared by centrifugation and protein concentrations were determined using the DC Protein Assay (BioRad) according to the manufacturer’s instructions. Equal amounts of protein for each sample (∼50 µg) were resolved by SDS-PAGE, transferred to a PVDF membrane, blocked in 5% nonfat dry milk (BioRad), and probed with either anti-phospho-p38 MAPK (Thr180/Tyr182; #9211, Cell Signaling Technology), anti-PMK-1 (Peterson et al. 2019), or anti-Actin antibodies (ab3280, Abcam).

For co-immunoprecipitation (co-IP) experiments, ∼50,000 day one adults were harvested, homogenized, and resuspended in lysis buffer exactly as previously described (Dowen 2019). Worm lysates were sonicated with the Bioruptor Pico sonication instrument (Diagenode) and cleared by centrifugation. For co-IP of mKate2::3xFLAG::DRL-1 with 3xHA::mGL::FLR-4 from strain DLS781, the lysate was split in three equal parts and subjected to a mock IP, anti-HA IP (2 µg of 3F10; 11867423001, Sigma), or anti-FLAG IP (5 µg of M2; F1804, Sigma). Prior to the immunoprecipitations antibodies were bound to Protein G Dynabeads (Invitrogen) with the mock IP sample lacking antibody. Western blotting of immunoprecipitated proteins was performed as described above using anti-FLAG (F1804, Sigma) or anti-HA (11867423001, Sigma) antibodies.

### EMS mutagenesis and identification of causative mutations

Mutagenesis of *drl-1(rhd109); mgIs70* (DLS364) animals with ethyl methanesulfonate (EMS, Sigma-Aldrich) was performed exactly as previously described (Dowen et al. 2016). A total of 50,000 haploid genomes were screened across 6 genetically distinct pools. Suppressor mutants (∼130 viable strains) were selected based on increased growth rate and/or *mgIs70[Pvit-3::GFP]* expression relative to the parental strain.

Suppressor strains were selected and backcrossed to the DLS364 strain two times. F2 recombinants from the second backcross that displayed the suppression phenotype were singled, the resulting plates were allowed to starve, the animals across all individual plates were pooled, and genomic DNA was prepared from the pool using the Qiagen Gentra Puregene Tissue Kit (Doitsidou et al. 2010). Whole genome sequencing libraries were prepared using the TruSeq DNA PCR-Free kit (Illumina) and sequenced on an Illumina HiSeq 4000 instrument according to the manufacturer’s instructions. Identification of candidate suppressor mutations was performed as previously described (Minevich et al. 2012) using in-house scripts.

### Reporter imaging and quantification

To measure P*vit-3::mCherry* fluorescence, strains carrying the *rhdSi42* reporter were grown to the L4 stage, picked to new plates, mounted 24 hours later as day 1 adults with 25mM levamisole on a 2% agarose pad, and imaged with a Nikon SMZ-18 Stereo microscope equipped with a DS-Qi2 monochrome camera. Worm bodies were traced in the brightfield channel and mean intensities (gray values) were calculated in the mCherry channel using ImageJ. The data were plotted in Prism 9 and a one-way ANOVA with a Bonferroni correction was performed to calculate *P* values.

For TIR-1::wrmScarlet imaging, animals were reared on *glo-3* RNAi for several generations using the RNAi-competent *E. coli* OP50(xu363) strain (Xiao et al. 2015). Knockdown of *glo-3* reduces intestinal autofluorescence and facilitates TIR-1 imaging (Peterson et al. 2022). At the L4 stage, animals were picked to fresh *glo-3* RNAi plates, mounted on a 2% agarose pad 24 hours later, and imaged with an CFI Apo 60X Oil TIRF objective on a Nikon Ti2 widefield microscope equipped with a Hamamatsu ORCA-Fusion BT camera. The *mKate2::drl-1*; *mGreenLantern::flr-4; glo-4(ok623)* CRISPR knock-in animals were also imaged at 60X on the Nikon Ti2 microscope.

TIR-1 puncta were measured in the posterior intestine since these 2-4 epithelial cells are not obstructed by other tissues. Z-stacks were cropped to a depth of 3-4 microns to capture planes where the intestine can be clearly visualized. All Z-stacks were denoised, deconvoluted, and compressed into a single image using the Nikon NIS-Elements analysis software. ROIs were hand drawn around the 2-4 posterior-most intestinal cells and the TIR-1 puncta were quantified using the object counts feature in NIS-Elements, which separated TIR-1::wrmScarlet puncta from the autofluorescent gut granules that appear in both the FITC and mCherry channels. TIR-1 puncta counts, puncta area, and puncta area relative to the ROI area were plotted in Prism 9 and a one-way ANOVA with a Bonferroni correction was performed to calculate *P* values.

For PHA-4::EGFP imaging, *reSi5; mKate2::3xFLAG::AID::drl-1; pha-4::EGFP* animals were reared on OP50(xu363) or HT115(DE3) *glo-3* RNAi bacteria in the presence of 4mM auxin for multiple generations to reduce gut autofluorescence and deplete the DLR-1 protein. L4 animals were picked to new plates and imaged 24 hours later as day 1 adults with a 10X objective on the Nikon Ti2 microscope. All Z-stacks were denoised and deconvoluted using the Nikon NIS-Elements analysis software and a single plane was selected for analysis based upon which had the best resolved intestinal nuclei. The nuclei of the two visible anterior and posterior-most intestinal cells were outlined by hand in ImageJ and GFP fluorescent signal intensity measurements were obtained by recording the mean gray value of each nucleus. The mean intensity values for each anterior nucleus for the auxin-treated animals (∼10 individuals/sample) were divided by the mean intensity value of all the anterior nuclei for the minus auxin control animals. An identical analysis was performed for the posterior nuclei and the data were plotted in Prism 9 and a one-way ANOVA with a Bonferroni correction was performed.

## ACKNOWLEDGEMENTS

The Caenorhabditis Genetics Center is supported by the NIH Office of Research Infrastructure Programs (P40 OD010440) and provided some of the strains used in this study. The total-PMK-1 antibody was generously provided by Dr. Read Pukkila-Worley (UMass). This work was supported by the Integrative Program for Biological and Genome Sciences (UNC) and NIGMS grant R35GM137985 to R.H.D.

## AUTHOR CONTRIBUTIONS

S.K.T, A.Y.P, and R.H.D designed the experiments and S.K.T, A.Y.P, P.C.B, N.R.C, and R.H.D performed the experiments and interpreted the results. S.K.T and R.H.D prepared the manuscript.

