## Supplemental Material for "Follicle stimulating hormone signaling opposes the DRL-1/FLR-4 MAP Kinases to balance p38-mediated growth and lipid homeostasis in *C. elegans*"

| **Contents** | **Page** |
| --- | --- |
| 1. Supplemental Tables S1-S5 | 2 |
| 2. Supplemental Figures S1-S9 | 9 |
| 3. References | 18 |

***drl-1(rhd109)* candidate suppresor mutations in DLS425**

| Chromosome | Position | Gene | Change |
|---|---|---|---|
| *II* | 2903191 | *Y110A2AM.1* | F270fs (Early stop) |
| *V* | 5916484 | *C04E6.11* | S358L |
| *V* | 7046939 | *F20A1.2* | G151R |
| *V* | 8159351 | *sul-2* | V395I |
| *V* | 9726592 | *aagr-3* | S849F (isoform a) |
| *V* | 9738884 | *aqp-4* | W106* (Stop) |
| *V* | 11749795 | *T04F3.1* | G2463E (isoform a) |
| ***V*** | **11937764** | ***flr-2*** | **R8* (Stop)** |
| *V* | 13415877 | *cut-2* | G197R |
| *V* | 13918750 | *str-250* | Y37F |
| *V* | 14923180 | *srg-54* | R60H |
| *V* | 16275184 | *T13F3.7* | K45R |

**Table S1. Identification of the *flr-2(rhd117)* mutation.** To identify the causative *drl-1(rhd109)* suppressor mutations, EMS mutants were backcrossed to DLS364, the independently segregating F2 animals displaying the suppression phenotypes were pooled, the genomic DNA was sequenced, and candidate mutations were identified as described in the Materials and Methods. The *flr-2* mutation (shown in bold) was selected for further analysis since it is predicted to be a strong loss-of-function allele. The resulting amino acid change is listed in the last column.

***drl-1(rhd109)* candidate suppresor mutations in DLS426**

| Chromosome | Position | Gene | Change |
|---|---|---|---|
| *II* | 2903191 | *Y110A2AM.1* | F270fs (Early stop) |
| *III* | 7363379 | *numr-1* | Y29S |
| *V* | 7329381 | *clec-7* | C273Y |
| *V* | 7599603 | *C50E3.9* | A234T |
| *V* | 7836320 | *C37C3.7* | G133E |
| *V* | 7982505 | *F26D11.1* | D132fs (Early stop) |
| *V* | 9261106 | *nhr-36* | E7K |
| ***V*** | **9889978** | ***fshr-1*** | **R7* (Stop)** |
| *V* | 10901577 | *F32D8.8* | S326F |
| *V* | 11442831 | *F57B7.2* | A151T (isoform a) |
| *V* | 11758540 | *T04F3.2* | A244V |
| *V* | 13897245 | *cyp-35C1* | T330S |
| *V* | 14326193 | *F47B8.5* | V207I (isoform a) |
| *V* | 14335951 | *srm-5* | T184I |
| *V* | 14855596 | *npr-13* | V288I |
| *V* | 15171024 | *srh-278* | Q161* (Stop) |
| *V* | 15720159 | *Y6E2A.7* | P78S |
| *V* | 16687549 | *srz-102* | T8I |
| *V* | 17034136 | *srz-94* | Splicing Variant |

**Table S2. Identification of the *fshr-1(rhd118)* mutation.** The candidate causative *drl-1(rhd109)* suppressor mutations were identified as described in Table S1 and the Materials and Methods. The *fshr-1* mutation (shown in bold) was selected for further analysis since it is predicted to be a strong loss-of-function allele. The resulting amino acid change is listed in the last column.

| Strain | Genotype | Reference |
|---|---|---|
| N2 | Wild-type | 1/7/23 9:27:00 PM |
| DLS362 | *drl-1(rhd109) IV* | This study |
| DLS363 | *drl-1(rhd110) IV* | This study |
| DLS364 | *drl-1(rhd109) IV; mgIs70[Pvit-3::GFP]* | This study |
| DLS365 | *drl-1(rhd110) IV; mgIs70[Pvit-3::GFP]* | This study |
| DLS425 | *drl-1(rhd109) IV; flr-2(rhd117) V; mgIs70[Pvit-3::GFP]* | This study |
| DLS426 | *drl-1(rhd109) IV; fshr-1(rhd118) V; mgIs70[Pvit-3::GFP]* | This study |
| DLS428 | *nhr-49(ok2165) I; drl-1(rhd109) IV* | This study |
| DLS513 | *drl-1(rhd109) IV; mgIs70[Pvit-3::GFP]; rhdEx99[Pvha-6::mCherry::his-58::SL2::drl-1 cDNA]* | This study |
| DLS514 | *drl-1(rhd109) IV; mgIs70[Pvit-3::GFP]; rhdEx100[Pvha-6::mCherry::his-58::SL2::drl-1 cDNA]* | This study |
| DLS515 | *drl-1(rhd109) IV; mgIs70[Pvit-3::GFP]; rhdEx101[Pcol-10::mCherry::his-58::SL2::drl-1 cDNA]* | This study |
| DLS516 | *drl-1(rhd109) IV; mgIs70[Pvit-3::GFP]; rhdEx102[Pcol-10::mCherry::his-58::SL2::drl-1 cDNA]* | This study |
| DLS519 | *drl-1(rhd109) IV; flr-2(ut5) V; mgIs70[Pvit-3::GFP]* | This study |
| DLS520 | *drl-1(rhd109) IV; fshr-1(ok778) V; mgIs70[Pvit-3::GFP]* | This study |
| DLS523 | *flr-4(ut7) X; mgIs70[Pvit-3::GFP]* | This study |
| DLS537 | *rhdSi42[Pvit-3::mCherry::unc-54 3'UTR + cb-unc-119(+)] II* | This study |
| DLS539 | *rhdSi42[Pvit-3::mCherry::unc-54 3'UTR + cb-unc-119(+)] alxIs9[Pvha-6::SID-1::SL2::GFP] II; sid-1(qt9) V* | This study |
| DLS581 | *drl-1(rhd175) IV; mgIs70[Pvit-3::GFP]* | This study |
| DLS582 | *drl-1(rhd176) IV; mgIs70[Pvit-3::GFP]* | This study |
| DLS583 | *drl-1(rhd177) IV; mgIs70[Pvit-3::GFP]* | This study |
| DLS586 | *drl-1(rhd180[P269S E270K]) IV; mgIs70[Pvit-3::GFP]* | This study |
| DLS622 | *reSi5[Pges-1::TIR1::F2A::mTagBFP2::NLS::AID::tbb-2 3'UTR] I; rhdSi42[Pvit-3::mCherry::unc-54 3'UTR + cb-unc-119(+)] II* | This study |
| DLS626 | *flr-4(n2259) X; mgIs70[Pvit-3::GFP]* | This study |
| DLS627 | *rhdSi42[Pvit-3::mCherry::unc-54 3'UTR + cb-unc-119(+)] II; fshr-1(rhd118) V* | This study |
| DLS628 | *drl-1(rhd197[E253A G254A P269S E270K]) IV; mgIs70[Pvit-3::GFP]* | This study |
| DLS632 | *drl-1(rhd197[E253A G254A P269S E270K]) IV* | This study |
| DLS634 | *rhdSi42[Pvit-3::mCherry::unc-54 3'UTR + cb-unc-119(+)] II; drl-1(rhd109) IV; fshr-1(rhd118) V* | This study |
| DLS635 | *rhdSi42[Pvit-3::mCherry::unc-54 3'UTR + cb-unc-119(+)] II; drl-1(rhd109) IV; flr-2(rhd117) V* | This study |
| DLS636 | *rhdSi42[Pvit-3::mCherry::unc-54 3'UTR + cb-unc-119(+)] II; drl-1(rhd109) IV* | This study |
| DLS637 | *rhdSi42[Pvit-3::mCherry::unc-54 3'UTR + cb-unc-119(+)] II; drl-1(rhd110) IV* | This study |
| DLS638 | *rhdSi42[Pvit-3::mCherry::unc-54 3'UTR + cb-unc-119(+)] II; flr-4(ut7) X* | This study |

| | | |
|---|---|---|
| DLS644 | *reSi5[Pges-1::TIR1::F2A::mTagBFP2::NLS::AID::tbb-2 3'UTR] I; rhdSi42[Pvit-3::mCherry::unc-54 3'UTR + cb-unc-119(+)] II; drl-1(rhd203[mKate2::TEV::3xFLAG::AID::drl-1]) IV* | This study |
| DLS657 | *rhdSi42[Pvit-3::mCherry::unc-54 3'UTR + cb-unc-119(+)] II; flr-2(rhd117) V; flr-4(ut7) X* | This study |
| DLS658 | *rhdSi42[Pvit-3::mCherry::unc-54 3'UTR + cb-unc-119(+)] II; fshr-1(rhd118) V; flr-4(ut7) X* | This study |
| DLS663 | *rhdSi42[Pvit-3::mCherry::unc-54 3'UTR + cb-unc-119(+)] II; flr-2(rhd117) V* | This study |
| DLS664 | *flr-2(rhd117) V* | This study |
| DLS665 | *drl-1(rhd109) IV; flr-2(rhd117) V* | This study |
| DLS674 | *reSi5[Pges-1::TIR1::F2A::mTagBFP2::NLS::AID::tbb-2 3'UTR] I; rhdSi42[Pvit-3::mCherry::unc-54 3'UTR + cb-unc-119(+)] II; flr-4(rhd209[mNG::TEV::3xFLAG::AID::flr-4]) X* | This study |
| DLS685 | *nhr-49(nr2041) I; rhdSi42[Pvit-3::mCherry::unc-54 3'UTR + cb-unc-119(+)] II; drl-1(rhd109) IV* | This study |
| DLS696 | *reSi7[Prgef-1::TIR1::F2A::mTagBFP2::AID::NLS::tbb-2 3'UTR] I; rhdSi42[Pvit-3::mCherry::unc-54 3'UTR + cb-unc-119(+)] II* | This study |
| DLS699 | *reSi7[Prgef-1::TIR1::F2A::mTagBFP2::AID::NLS::tbb-2 3'UTR] I; rhdSi42[Pvit-3::mCherry::unc-54 3'UTR + cb-unc-119(+)] II; flr-4(rhd209[mNG::TEV::3xFLAG::AID::flr-4]) X* | This study |
| DLS700 | *reSi1[Pcol-10::TIR1::F2A::mTagBFP2::AID::NLS::tbb-2 3'UTR] I; rhdSi42[Pvit-3::mCherry::unc-54 3'UTR + cb-unc-119(+)] II; drl-1(rhd203[mKate2::TEV::3xFLAG::AID::drl-1]) IV* | This study |
| DLS701 | *reSi1[Pcol-10::TIR1::F2A::mTagBFP2::AID::NLS::tbb-2 3'UTR] I; rhdSi42[Pvit-3::mCherry::unc-54 3'UTR + cb-unc-119(+)] II* | This study |
| DLS712 | *rhdSi42[Pvit-3::mCherry::unc-54 3'UTR + cb-unc-119(+)] alxIs9[Pvha-6::sid-1::SL2::GFP] II; drl-1(rhd109) IV; sid-1(qt9) V* | This study |
| DLS781 | *drl-1(rhd203[mKate2::TEV::3xFLAG::AID::drl-1]) IV; glo-4(ok623) V; flr-4(rhd244[3xHA::mGL::flr-4]) X* | This study |
| DLS831 | *reSi5[Pges-1::TIR1::F2A::mTagBFP2::NLS::AID::tbb-2 3'UTR] I; drl-1(rhd203[mKate2::TEV::3xFLAG::AID::drl-1]) IV; pha-4(st12220[pha-4::TY1::EGFP::3xFLAG]) V* | This study |
| DLS835 | *tir-1(ums63[tir-1::wrmScarlet]) III; drl-1(rhd109) IV; flr-2(rhd117) V* | This study |
| DLS836 | *reSi5[ges-1p::TIR1::F2A::mTagBFP2::AID*::NLS::tbb-2 3'UTR] I; drl-1(rhd203[mKate2::TEV::3xFLAG::AID::drl-1]) pmk-1(km25) IV* | This study |
| DLS837 | *rhdSi46[Psng-1::mCherry::his-58::SL2::flr-2 + cb-unc-119(+)] II; drl-1(rhd109) IV; flr-2(rhd117) V* | This study |
| DLS841 | *tir-1(ums63[tir-1::wrmScarlet]) III; drl-1(rhd109) IV* | This study |
| DLS845 | *reSi5[ges-1p::TIR1::F2A::mTagBFP2::AID*::NLS::tbb-2 3'UTR] I; drl-1(rhd203[mKate2::TEV::3xFLAG::AID::drl-1]) pmk-1(km25) IV; pha-4(st12220) V* | This study |
| DLS846 | *reSi5[ges-1p::TIR1::F2A::mTagBFP2::AID*::NLS::tbb-2 3'UTR] I; drl-1(rhd203[mKate2::TEV::3xFLAG::AID::drl-1])* | This study |

| | | |
|---|---|---|
| DLS848 | *reSi5[ges-1p::TIR1::F2A::mTagBFP2::AID*::NLS::tbb-2 3'UTR] I; drl-1(rhd203[mKate2::TEV::3xFLAG::AID::drl-1]) IV; flr-2(rhd117) V* | This study |
| GR2122 | *mgIs70[Pvit-3::GFP]* | (Dowen et al. 2016) |
| IG544 | *nipi-3(fr4) X* | (Pujol et al. 2008) |
| JC2209 | *olrn-1(ut305) X* | (Torayama et al. 2007) |
| JC49 | *flr-2(ut5) V* | (Take-Uchi et al. 1998) |
| JC51 | *flr-4(ut7) X* | (Take-uchi et al. 2005) |
| KG421 | *gsa-1(ce81)* | (Schade et al. 2005) |
| KG532 | *kin-2(ce179)* | (Schade et al. 2005) |
| MGH171 | *sid-1(qt9) V; alxIs9[Pvha-6::SID-1::SL2::GFP]* | (Melo and Ruvkun 2012) |
| MT5701 | *flr-4(n2259) X* | (Take-uchi et al. 2005) |
| QK52 | *rde-1(ne219) V; xkIs99[Pwrt-2::rde-1]* | (Melo and Ruvkun 2012) |
| RB911 | *fshr-1(ok778) V* | (Cho et al. 2007; C. elegans Deletion Mutant Consortium 2012) |
| RPW403 | *tir-1(ums63[tir-1::wrmScarlet]) III* | (Peterson et al. 2022) |
| RPW43 | *nsy-1(ums8) II; agIs44[pF08G5.6::GFP::unc-54(3'UTR) + Pmyo-2::mCherry]* | (Cheesman et al. 2016) |

**Table S3. *C. elegans* strains used in this study.** The strain names, genotypes, and any associated references are shown.

| Target Gene | Location in Gene | crRNA Sequence | Alleles Generated |
|---|---|---|---|
| *drl-1* | 5' end | UCCGUCAAAAAUGCAUUCAGGUUUUAGAGCUAUGCU | *rhd203* |
| *drl-1* | Internal | UCUAAUGACCGGAACGCUUCGUUUUAGAGCUAUGCU | *rhd180* |
| *drl-1* | Internal | CUGGCGGAUCCUUUUAUUGAGUUUUAGAGCUAUGCU | *rhd197* |
| *flr-4* | 5' end | UAAUUUAUUGGCAUUCCCGUGUUUUAGAGCUAUGCU | *rhd209, rhd244* |

**Table S4. The crRNAs used in this study.** A list of the crRNA guides, including their genomic targets and ribonucleotide sequences, that were used in this study. The alleles generated using CRISPR/Cas9 gene editing are also shown (far right column).

| mRNA Target | Primer Sequence (5' to 3') | Reference |
|---|---|---|
| *act-1* | F: GCTGGACGTGATCTTACTGATTACC<br>R: GTAGCAGAGCTTCTCCTTGATGTC | (Hoogewijs et al. 2008) |
| *vit-1* | F: GAGGTTCGCTTTGACGGATA<br>R: GGCTTCACATTCCTCGTTCT | (Ding and Grosshans 2009) |
| *vit-2* | F: GACACCGAGCTCATCCGCCCA<br>R: TTCCTTCTCTCCATTGACCT | (DePina et al. 2011) |
| *vit-3/4/5* | F: CATGTGCACCATCGAAGAACTC<br>R: CCAATGTGGTTTCAATGACAAGTTG | (Dowen et al. 2016) |
| *vit-6* | F: TTCACCCAGAAGCCAGTTC<br>R: AGGATGGGAGGCAGTAGAC | (Dowen et al. 2016) |
| *ech-9* | F: AGGAAAATGGACTTGAGCCG<br>R: CTTTCCGTTGGGTTTTATCGTC | This study |
| *ugt-18* | F: AACCGGCACTGATAATTCCCCTTATGG<br>R: TAGAGCCCCATGTTCAACTGC | (Chamoli et al. 2014) |
| *tir-1* | F: GGGTTCACGACTATCAGGATG<br>R: TTCGGGAGATAGAAGGCATTTC | This study |
| *nsy-1* | F: CAGAACAACAAGAGGCAAGTG<br>R: GAAGTTGCAGCCATTCGAC | This study |
| *sek-1* | F: CACTGTTTGGCGACGATGAG<br>R: ATTCCGTCCACGTTGCTGAT | (Wu et al. 2021) |
| *pmk-1* | F: TTGATGTATGGTCAGTTGGG<br>R: GATCGATGTGATCAGATCCAG | (Amrit et al. 2019) |
| *vhp-1* | F: CCTCTCACAACCATGTCTATCC<br>R: CATGGTCTCATCCAAGCTATCA | (Wu et al. 2021) |
| *cebp-1* | F: GCAAGACAAGACTCTCTTAC<br>R: CCAAGGTCCAGCTCAGTTTC | (Wu et al. 2021) |

**Table S5. The RT-qPCR primers.** The primer sequences (5' to 3') and any associated references are shown for the qPCR primers used in this study.

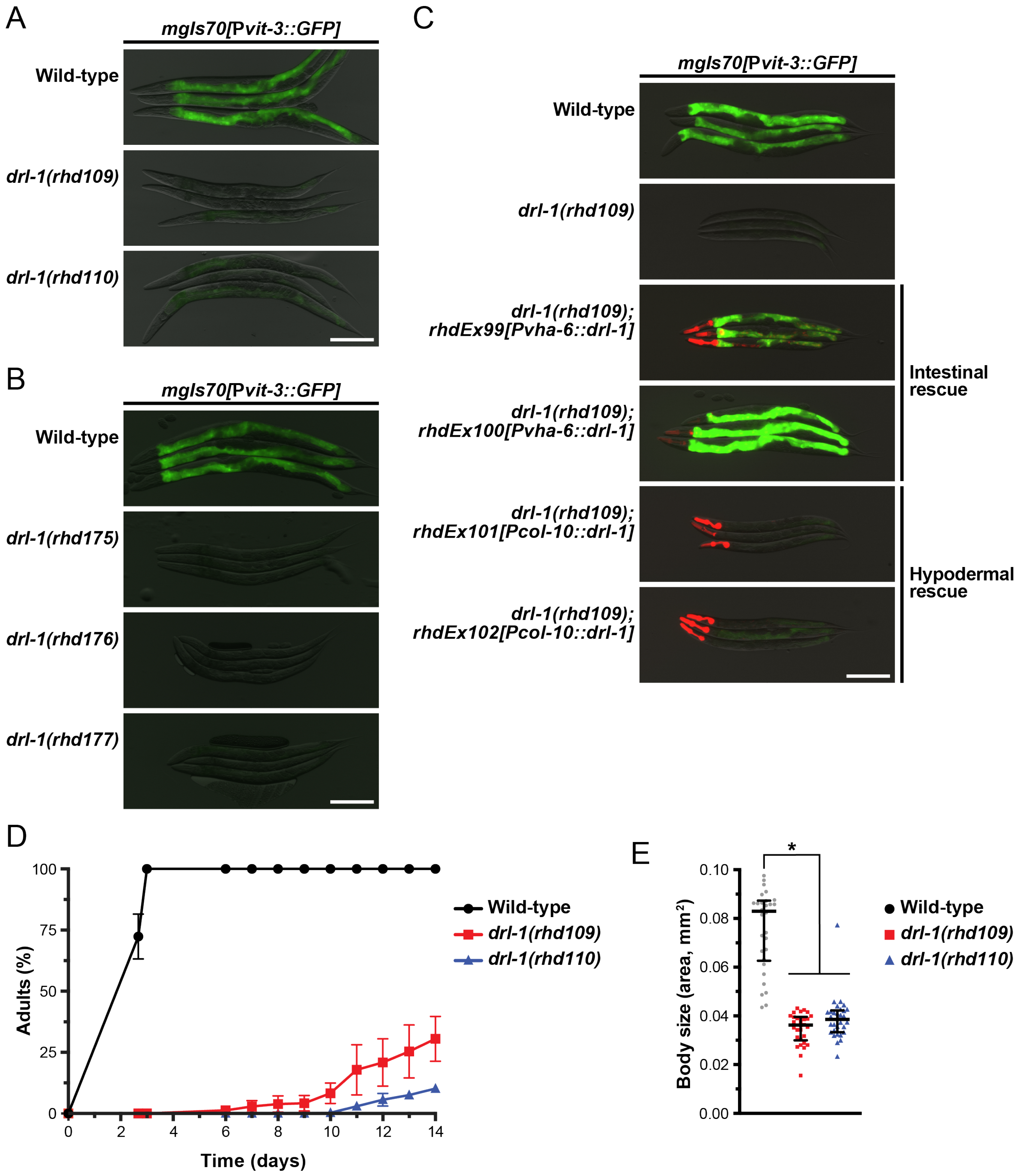


**Figure S1. Intestinal DRL-1 controls vitellogenin production and growth.** (A, B) Representative fluorescence and DIC overlaid images of *mgIs70[*P*vit-3::GFP]* reporter expression in various *drl-1* mutants (scale bar, 200 µm). The *mgIs70* transgene is a high-copy transgene. (C) Intestinal, but not hypodermal, rescue of *drl-1(rhd109)* mutants with *drl-1* cDNA restores P*vit-3::GFP* expression (scale bar, 200 µm). (D) Growth rate (mean +/- SEM) and (E) body size (day 1 adults; median and interquartile range; *, *P*<0.0001, one-way ANOVA) of wild-type and *drl-1* mutant animals.

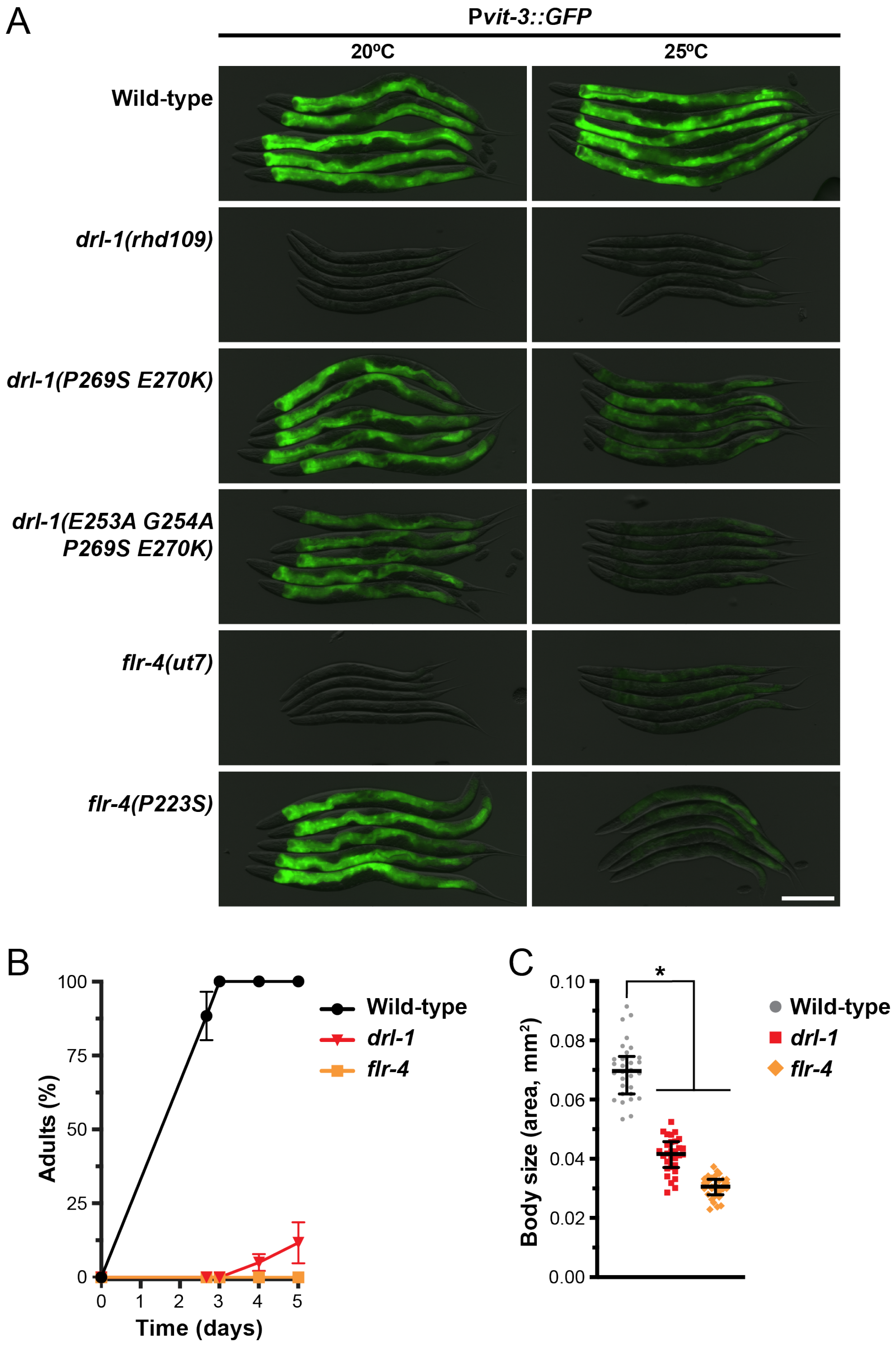


**Figure S2.** ***drl-1*** **and** ***flr-4*** **mutants display similar vitellogenesis and growth defects.** (A) Representative overlaid DIC and GFP fluorescence images of day 1 adult wild-type and mutant animals reared at 20 or 25°C (scale bar, 200 µm). (B) Growth rate (mean +/- SEM) and (C) body size (day 1 adults; median and interquartile range; *, $P$<0.0001, one-way ANOVA) of wild-type, *drl-1(rhd109)*, and *flr-4(ut7)* animals.

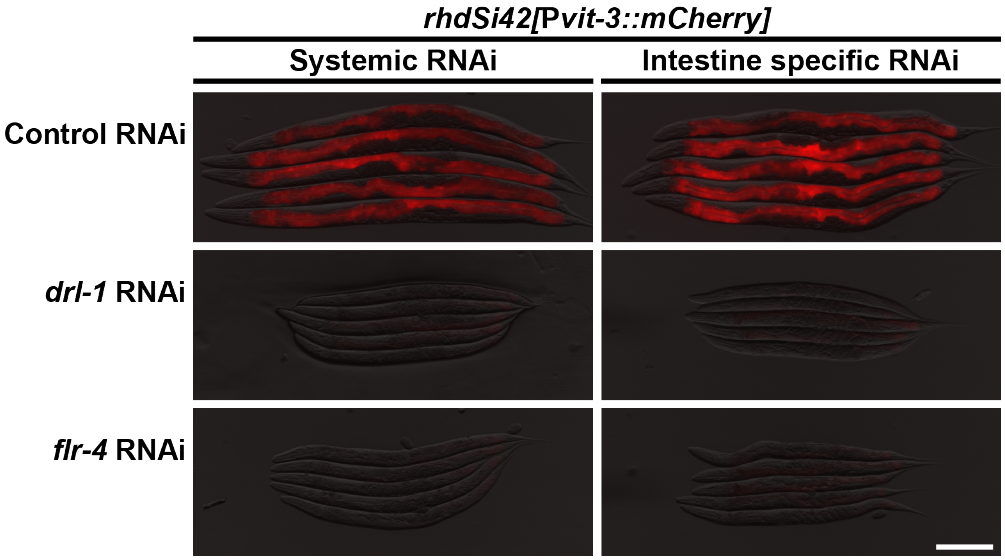


**Figure S3. *drl-1* and *flr-4* function in a cell-autonomous manner to regulate vitellogenin production.** Representative fluorescence images of P*vit-3::mCherry* reporter expression in day 1 adult animals after whole-body or tissue-specific knockdown of *drl-1* or *flr-4* by RNAi (scale bar, 200 µm).

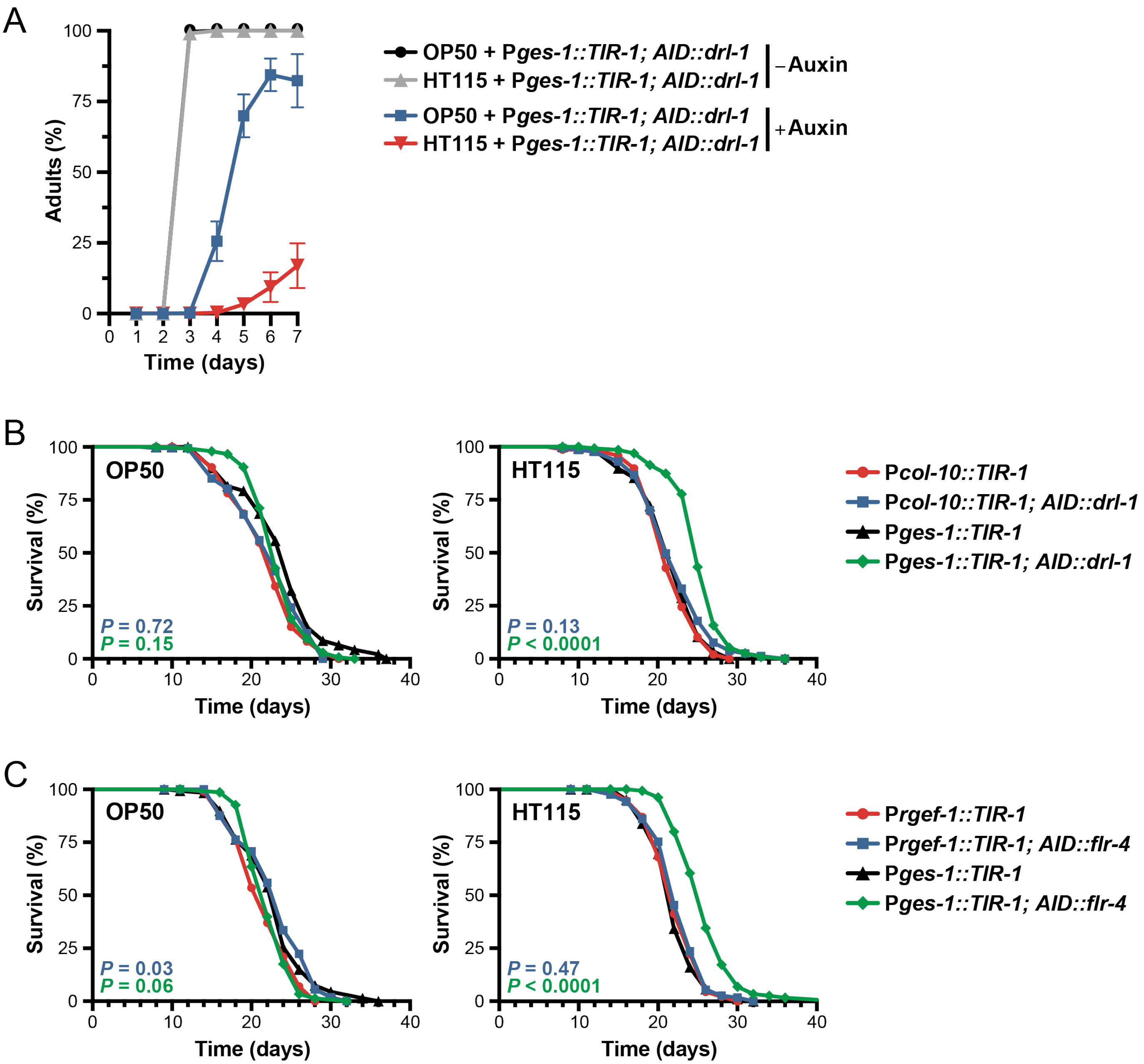


**Figure S4. Intestinal depletion of DRL-1 or FLR-4 paired with an *E. coli* HT115 diet results in severe growth defects and an extended lifespan.** (A) Growth rate (mean +/- SEM) of *mKate2::3xFLAG::AID::drl-1* animals with or without 4 mM auxin reared on *E. coli* OP50 or HT115. Longitudinal lifespan assays of (B) *mKate2::3xFLAG::AID::drl-1* or (C) *mNG::3xFLAG::AID::flr-4* animals grown at 20°C with FUDR on *E. coli* OP50 or HT115 (P*ges-1::TIR1*, intestinal depletion; P*col-10::TIR1*, hypodermal depletion; P*rgef-1::TIR1*, pan-neuronal depletion). Control animals only carry the *TIR1* transgenes and all strains were reared on 4 mM auxin from hatching.

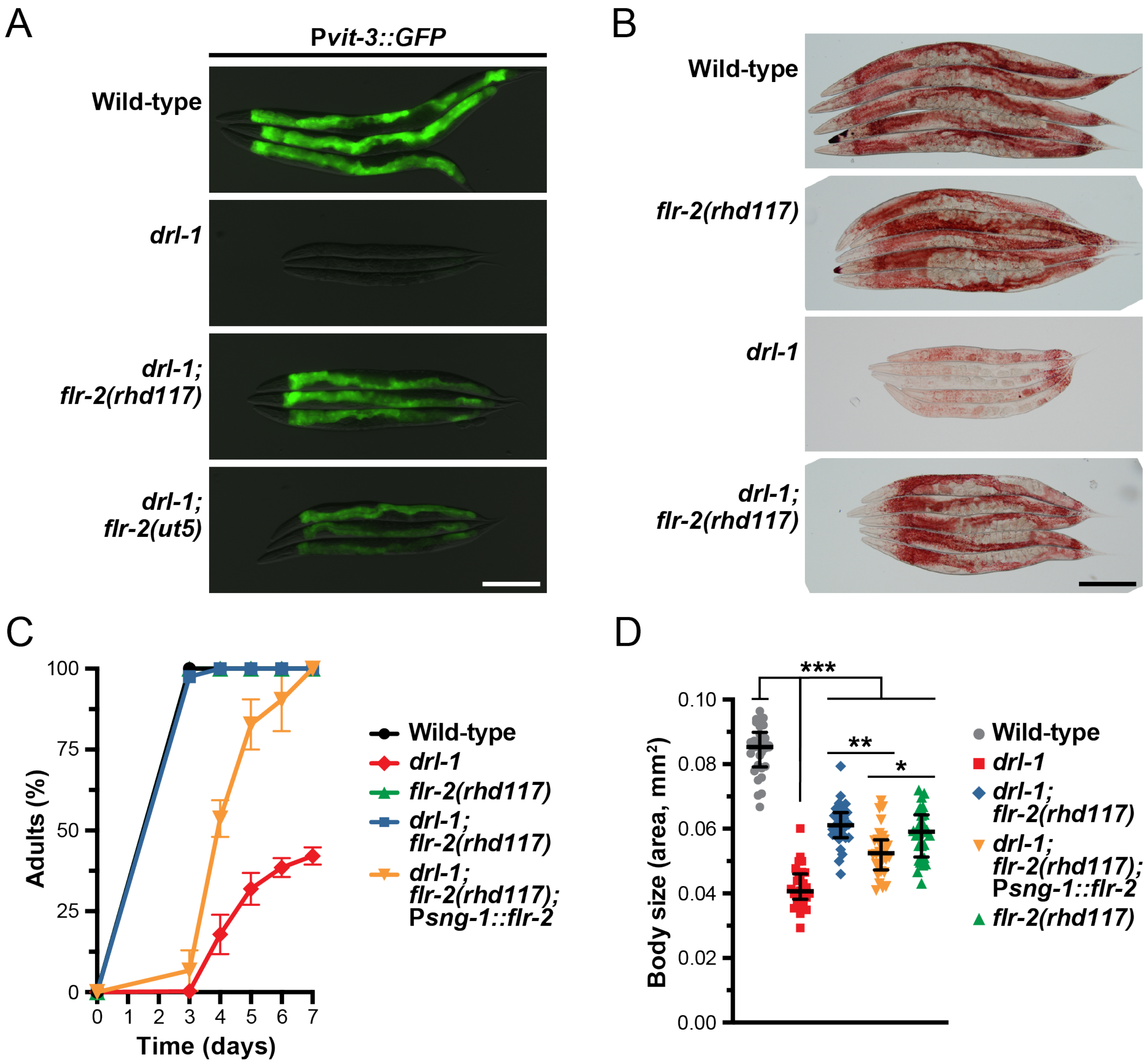


**Figure S5.** ***flr-2*** **functions in a non-cell-autonomous manner to suppress the** ***drl-1(rhd109)*** **mutation.** (A) Overlaid DIC and GFP fluorescence images of day 1 adult wild-type, *drl-1(rhd109)*, and *drl-1(rhd109)* double mutant animals (scale bar, 200 µm). (B) Representative images of day 1 adult wild-type, *flr-2(rhd117)*, *drl-1(rhd109)*, and *drl-1(rhd109); flr-2(rhd117)* animals stained with Oil Red O (scale bar, 200 µm). (C) Growth rate (mean +/- SEM) and (D) body size (day 1 adults) of wild-type, *drl-1(rhd109)* single and double mutants, and *flr-2* pan-neuronal rescue animals (P*sng-1::flr-2* is a single-copy rescue transgene). (D) Body size data are presented as the median and interquartile range (***, $P<0.0001$, **, $P=0.0003$, *, $P=0.04$, one-way ANOVA). Wild-type animals have a significantly larger body size compared to all other strains ($P<0.0001$).

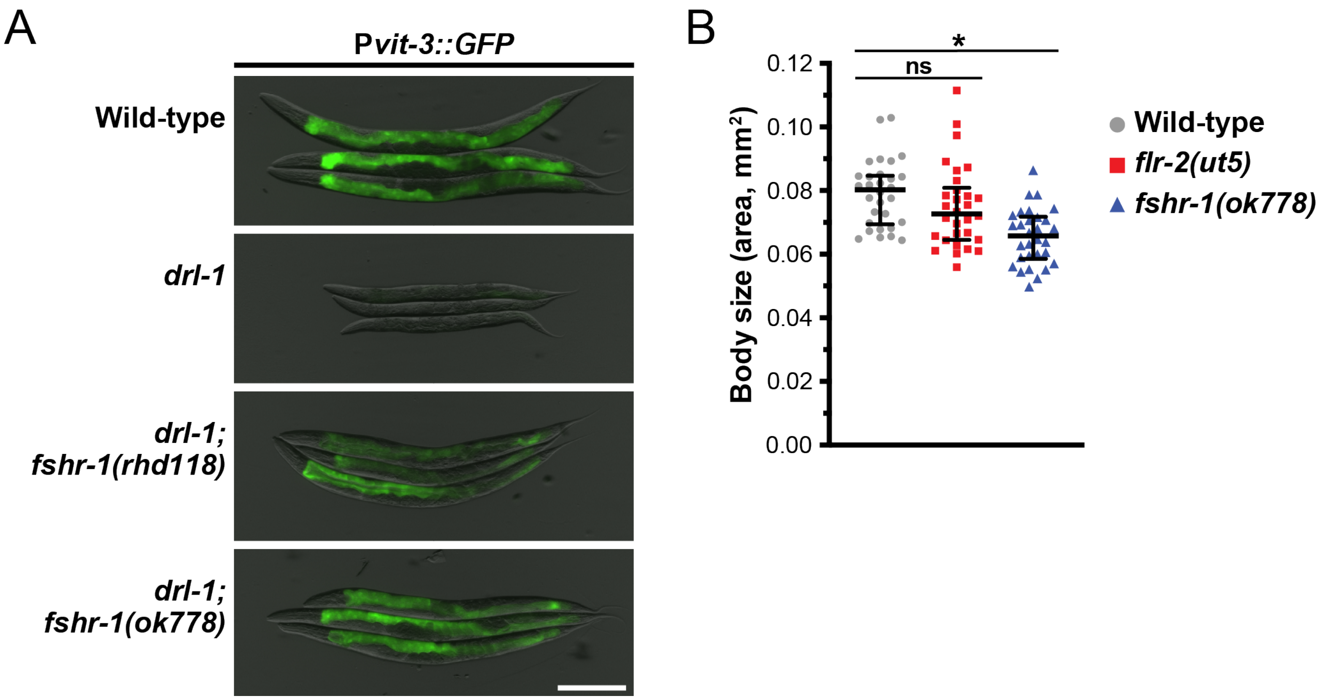


**Figure S6.** ***fshr-1*** **mutations suppress the vitellogenin expression defects displayed by the** ***drl-1(rhd109)*** **mutant.** (A) Representative overlaid DIC and GFP fluorescence images of day 1 adult wild-type, *drl-1(rhd109)*, and *drl-1(rhd109); fshr-1* double mutant animals (scale bar, 200 µm). (B) Body size of day 1 adult wild-type, *flr-2(ut5)*, and *fshr-1(ok778)* animals showing that both mutants are modestly smaller than wild-type (*, $P$<0.0001, ns, not significant, one-way ANOVA).

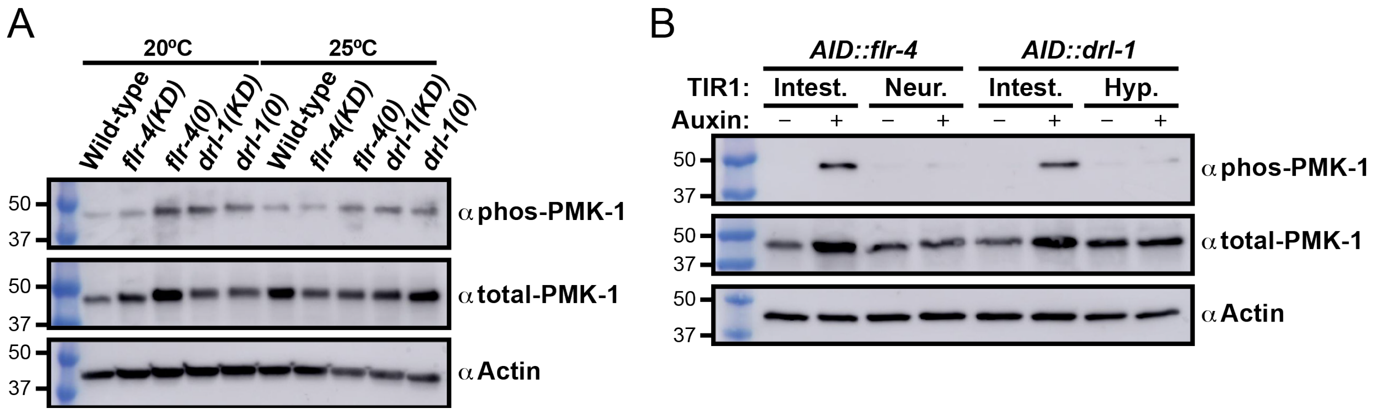


**Figure S7. Loss of intestinal DRL-1/FLR-4 hyperactivates the p38/PMK-1 signaling pathway.** (A) Western blot analysis of phospho-PMK-1, total-PMK-1, and actin levels in wild-type, kinase dead (KD) mutants, and null (0) mutants reared at 20 or 25°C. (B) Western blot analysis of phospho-PMK-1, total-PMK-1, and actin levels in *AID::drl-1* or *AID::flr-4* animals grown with or without 4 mM auxin (Intest., intestinal depletion, P*ges-1::TIR1*; Hyp., hypodermal depletion, P*col-10::TIR1*; Neur., pan-neuronal depletion; P*rgef-1::TIR1*). The *AID::drl-1* and *AID::flr-4* strains contain the *rhdSi42* transgene.

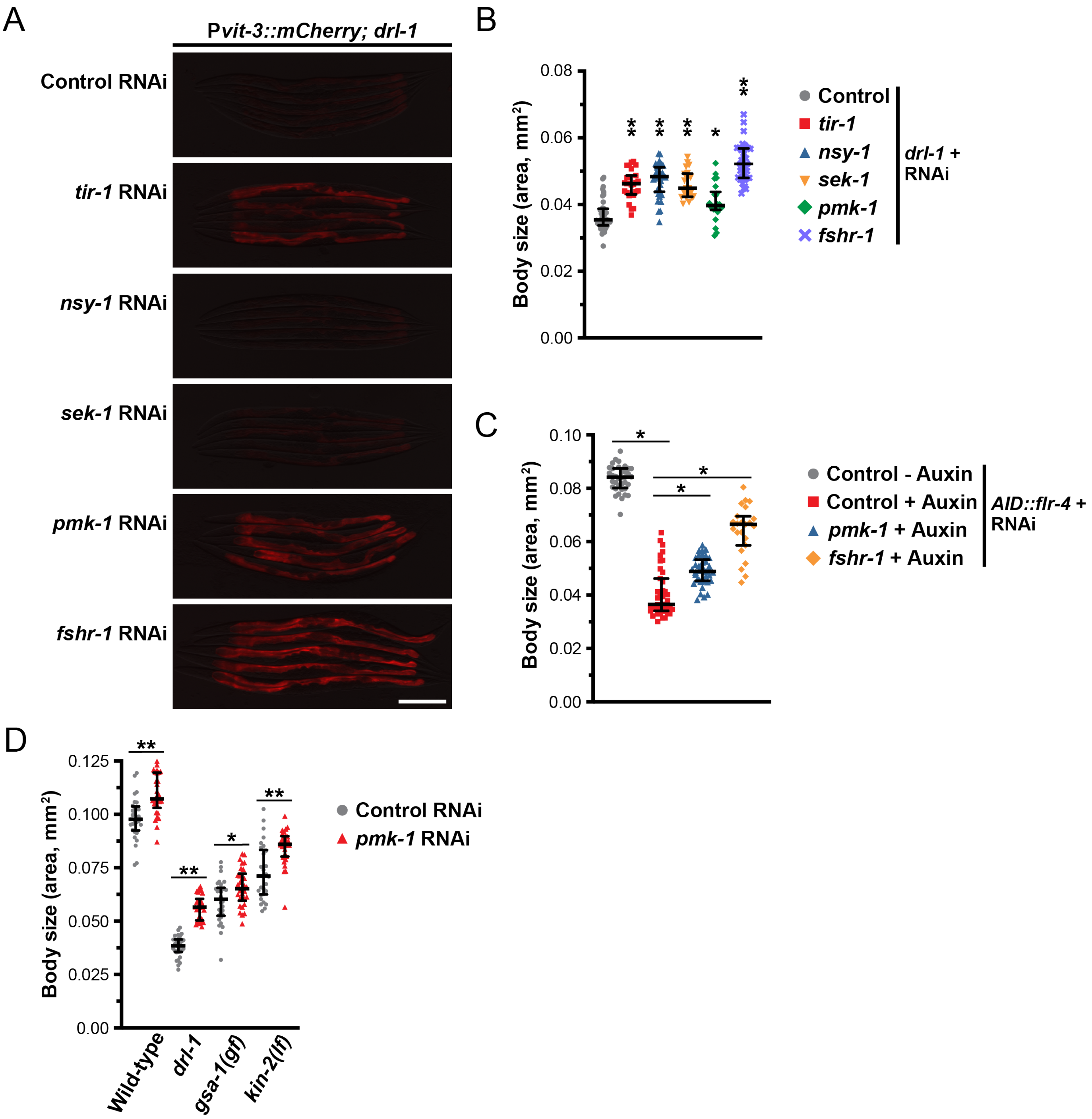


**Figure S8. Reduced p38/PMK-1 signaling suppresses the effects of losing DRL-1/FLR-4.** (A) Representative overlaid DIC and mCherry fluorescence images (scale bar, 200 µm) and (B) body size (median and interquartile range; *, *P*<0.02, **, *P*<0.001, one-way ANOVA) of day 1 adult *drl-1(rhd109)* animals after knock-down p38/PMK-1 pathway components by RNAi. (C) Body size of P*ges-1::TIR1*; *mNG::3xFLAG::AID::flr-4* animals after simultaneous depletion of intestinal AID::FLR-4 (with 4 mM auxin) and knock-down of *pmk-1* or *fshr-1* by RNAi. Data are shown as the median and interquartile range (*, *P*<0.0001, one-way ANOVA). (D) Body size of wild-type and the indicated mutants after control or *pmk-1* RNAi. These mutations have been previously shown to activate PKA signaling (lf, loss-of-function; gf, gain-of-function).

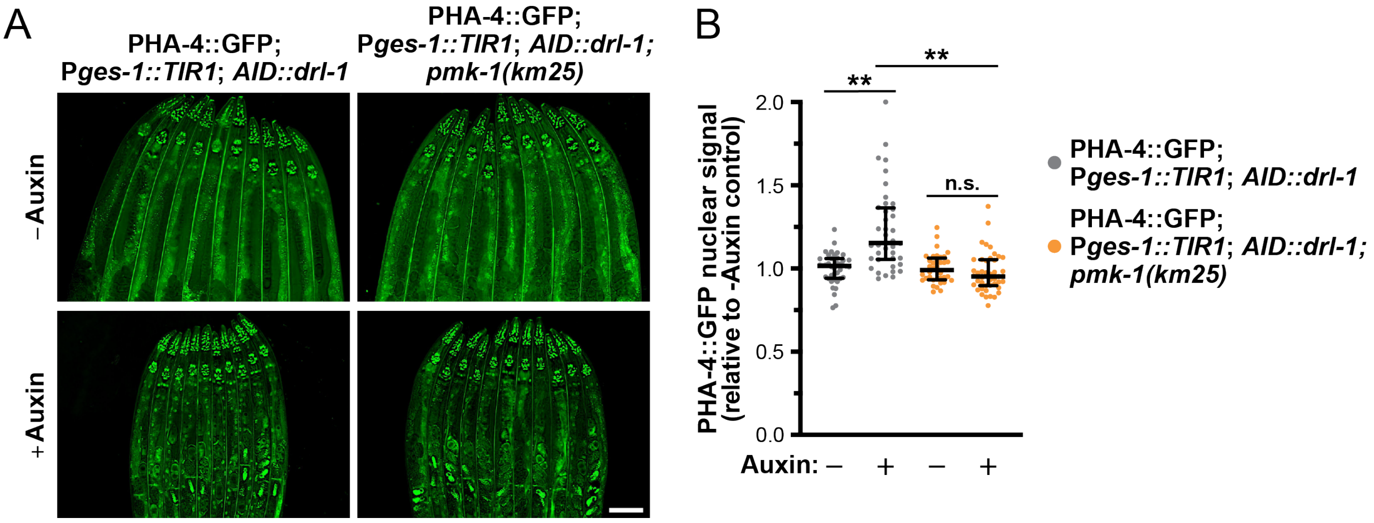


**Figure S9. DRL-1 inhibits the accumulation of nuclear PHA-4 through p38/PMK-1 signaling irrespective of diet.** (A) Fluorescence images (scale bar, 100 µm) and (B) quantification (median and interquartile range; n.s., not significant, **, $P$<0.0001, one-way ANOVA) of PHA-4::GFP nuclear localization after depletion of intestinal AID::DRL-1 using 4 mM auxin in wild-type or *pmk-1(km25)* animals grown on *E. coli* HT1115.
